# Profiling the Human Phosphoproteome to Estimate the True Extent of Protein Phosphorylation

**DOI:** 10.1101/2021.04.14.439901

**Authors:** Anton Kalyuzhnyy, Patrick A. Eyers, Claire E. Eyers, Zhi Sun, Eric W. Deutsch, Andrew R. Jones

## Abstract

Mass spectrometry-based phosphoproteomics allows large-scale generation of phosphorylation site data. However, analytical pipelines need to be carefully designed and optimised to minimise incorrect identification of phosphopeptide sequences or wrong localisation of phosphorylation sites within those peptides. Public databases such as PhosphoSitePlus (PSP) and PeptideAtlas (PA) compile results from published papers or openly available MS data, but to our knowledge, there is no database-level control for false discovery of sites, subsequently leading to the likely overestimation of true phosphosites. It is therefore difficult for researchers to assess which phosphosites are “real” and which are likely to be artefacts of data processing. By profiling the human phosphoproteome, we aimed to estimate the false discovery rate (FDR) of phosphosites based on available evidence in PSP and/or PA and predict a more realistic count of true phosphosites. We ranked sites into phosphorylation likelihood sets based on layers of accumulated evidence and then analysed them in terms of amino acid conservation across 100 species, sequence properties and functional annotations of associated proteins. We demonstrated significant differences between the sets and developed a method for independent phosphosite FDR estimation. Remarkably, we estimated a false discovery rate of 86.1%, 95.4% and 82.2% within sets of described phosphoserine (pSer), phosphothreonine (pThr) and phosphotyrosine (pTyr) sites respectively for which only a single piece of identification evidence is available (the vast majority of sites in PSP). Overall, we estimate that ∼56,000 Ser, 10,000 Thr and 12,000 Tyr phosphosites in the human proteome have truly been identified to date, based on evidence in PSP and/or PA, which is lower than most published estimates. Furthermore, our analysis estimated ∼91,000 Ser, 49,000 Thr and 26,000 Tyr sites that are likely to represent false-positive phosphosite identifications. We conclude that researchers should be aware of the significant potential for false positive sites to be present in public databases and should evaluate the evidence behind the phosphosites used in their research.

## Introduction

Protein phosphorylation is a fundamental post-translation modification (PTM) which regulates protein function and is very well-studied in relation to cell signalling pathways and disease (Cohen, 2001; Cohen, 2002; Goedert *et al*., 1992). Huge numbers of phosphorylated peptides and sites have been reported and characterized after isolation from human cells using approaches allied to tandem mass spectrometry (LC-MS/MS) which have primarily focussed on the phosphorylation of serine (Ser), threonine (Thr) and tyrosine (Tyr) residues (Amanchy *et al*., 2005; Nousiainen *et al*., 2006; Olsen *et al*., 2006; Olsen *et al*., 2010; Sharma *et al*., 2014). However, large numbers of non-canonical phosphorylation sites have also been annotated on proteins fractionated from human cells (Hardman *et al*., 2019). This additional complexity highlights the ongoing requirement for careful, evidence-based phosphosite identification from mass spectrometric datasets.

Historically, the analysis of phosphorylation sites in proteins tended to be accurate, initially relying on chromatography and solid-state Edman sequencing (Alessi *et al*., 1996; Wettenhall, Aebersold and Hood, 1991; Wettenhall, Erikson and Maller, 1992). However, such low-throughput approaches are now very rare, lacking the depth of coverage needed in modern proteomic workflows. The dominance of MS approaches has led to the development of multiple strategies to both understand and help mitigate the high levels of phosphopeptide false discovery rate (FDR), particularly in sets of mapped peptide spectral matches (PSMs) that result from LC-MS/MS and sequence database analysis (Elias and Gygi, 2007; Käll, Storey and Noble, 2009; Söderholm *et al*., 2014). The goal of such approaches is to separate true identifications from false ones. Even without considering non-canonical phosphorylation (which is likely not detectable in standard phosphoproteomics pipelines), many confidently-identified phosphopeptides possess multiple Ser, Thr or Tyr residues that could explain the number of phosphosites in a given proteolytically-generated peptide (Hardman *et al*., 2019). As such, statistical processes are applied, either within the search engine used for peptide mapping, or in a downstream software application to calculate additional statistics, such as a local *false localisation rate* (FLR) or conversely the probability that a given site within a peptide is correct or incorrect. Software / algorithms include phosphoRS (Taus *et al*., 2011), Ascore (Beausoleil *et al*., 2006), Andromeda’s PTM Score (Cox *et al*., 2011) and recently released PTMProphet (Shteynberg *et al*., 2019). We have previously benchmarked performance of some of the instruments and software pipelines for phosphoproteomics (Ferries *et al*., 2017), demonstrating that there is considerable variability in how such scores map to robust statistics, such as local or global FLR, depending on instrument fragmentation mode and resolution.

Following confident identification of phosphopeptides and localisation of given sites, data tend to be compiled from within a single study or across multiple studies (meta-analysis) to determine the extent of evidence for a given site from multiple PSMs. In general, where there are independent observations of PSMs supporting a phosphosite, it can be reasonably assumed that the evidence for a site to be real increases, although to our knowledge there are no current statistical models to calculate this phenomenon accurately. Multiple PSMs can be observed per identified phosphosite as a result of either different peptide sequences containing that site, or the same peptide sequence being detected several times (Dephoure *et al*., 2013). There are some caveats to this logic though, as it is possible for the same PSM to be wrongly assigned to a phosphopeptide multiple times. This can occur if the correct interpretation for the spectrum had a very similar peptide sequence and identical mass to the wrongly assigned phosphopeptide (Chalkley and Clauser, 2012; Lee, Jones and Hubbard, 2015). Although LC-MS/MS and computational analysis is generally recognised as very effective and reliable for phosphosite detection, from each study it is likely that there is some element of remaining false discovery of peptides and sites wrongly localised, depending on the statistical thresholds applied. This is particularly problematic for studies that set relatively weak thresholds (e.g., equating to site probability >0.75) in order to maximise sensitivity – more true positives may be gained, but at the expense of very large numbers of false positives passing the threshold. Methods and guidelines for FLR are still evolving and not consistently applied in published phosphoproteome studies, and so it is likely that most published studies contain considerable numbers of falsely localised phosphosites (Desiere *et al*., 2006; Dinkel *et al*., 2011; Gnad, Gunawardena and Mann, 2011; Hornbeck *et al*., 2015). This can lead to overestimation of the total number of known true human phosphosites if database providers do not control for FDR across multiple data sets (Ochoa *et al*., 2020).

One such database is PhosphoSitePlus (PSP) which represents a comprehensive, manually-curated and well-cited resource containing experimentally defined PTMs primarily focusing on phosphorylation (Hornbeck *et al*., 2015). As of March 2020, PSP encompassed phosphosite identification evidence across 17830 human protein sequences. The evidence for phosphorylation comes from manually-curated reviews of literature describing tandem MS and low-throughput experiments (Hornbeck *et al*., 2015). Interestingly, the majority of phosphosites in PSP only have 1 piece of evidence associated with their identification, and, as mentioned in the PSP documentation itself, the researchers must be cautious when accepting such sites as true positives (Hornbeck *et al*., 2015). It is possible that many users of PSP are not aware of the need for caution when reviewing or re-using data downloaded in bulk, and we are not aware of any previous effort to assess phosphosite FDR within PSP. A second curated proteomics resource is PeptideAtlas (PA) (Desiere *et al*., 2006) which is a repository of tandem MS datasets that have been processed through Trans-Proteomic Pipeline to ensure high and consistent quality of phosphopeptide identifications (Deutsch *et al*., 2010). The latest PA builds incorporate the use of the PTMProphet algorithm for phosphosite localisation where each potential phosphosite within an observed PSM is assigned a probability score between 0 and 1 of being phosphorylated (Shteynberg *et al*., 2019). As with PSP datasets, researchers must be cautious when accepting sites in PA with only 1 PSM as positively identified phosphosites. Instead, phosphosites that not only have multiple PSM observations in PA, but also have high phosphorylation probability scores assigned within the majority of those PSMs are most likely to be true positive identifications.

The issue of FDR within human phosphorylation databases is undeniably important, although to our knowledge no estimates have been made to predict its scale across large datasets. In this work, by profiling the human phosphoproteome, we aimed to estimate the false discovery rate of phosphosites with evidence in PSP and/or PA and use these estimates to predict the count of true phosphosites within the proteome. We categorised the sites into sets of various predicted phosphorylation likelihood based on the amount of positive identification evidence they have in PSP and PA. By using orthogonal properties of phosphosites assigned to the sets, such as evolutionary conservation, sequence properties and functional annotations, we aimed to demonstrate significant differences between the sets and develop an improved method for independent FDR estimation which can be used to indicate the extent of true phosphosites within the human phosphoproteome.

## Methods

### Processing and categorising phosphorylation data in PSP and PA

Phosphorylation data in PeptideAtlas (PA) (2020 build) (Desiere *et al*., 2006) was filtered to only include human Ser/Thr/Tyr sites from canonical UniProt protein sequences with at least 1 PSM observation (1,069,709 sites across 63,616 sequences) (SI Table 1). The sites were categorised according to the number of PSM observations with a certain phosphorylation probability score assigned by PTMProphet (Shteynberg *et al*., 2019). The counts of observations with a probability of >0.95 were used as “positive” evidence for site phosphorylation. The counts at a probability threshold of ≤ 0.19 were used as “negative” evidence in favour of a site being a non-phosphosite. The total number of PSM observations per site was considered to distinguish sites for which ≥10% of all associated PSMs had a PTM probability >0.95, from sites where a small minority (<10%) of associated PSMs had this probability. Based on this, selected confidence categories were applied to predict site phosphorylation likelihood in PA (“*High*”: ≥5 positive observations which accounted for ≥10% of total observations across all probabilities; “*Medium*”: ≥5 positive observations which accounted for <10% of total observations or 2-4 positive observations; “*Low*”: 1 positive observation; “*Not phosphorylated*”: 0 positive observations and ≥5 negative observations; “*Other*”: site did not fall into any described categories). PhosphoSitePlus (PSP) data (11/03/20 build; Phosphorylation_site_dataset.gz) (Hornbeck *et al*., 2015) was filtered to only include human Ser/Thr/Tyr sites from canonical protein sequences labelled by UniProt identifiers (231,607 sites across 17,830 sequences) (SI Table 2). The sites were ranked based on the number of times they have been characterised in low/high throughput studies. The sum of observations across all studies was used to predict site phosphorylation likelihood in PSP (“*High*”: ≥5 observations; “*Medium*”: 2-4 observations; “*Low*”: 1 observation).

**Table 1.** Categorising serine (Ser), threonine (Thr) and tyrosine (Tyr) sites from UniProt’s reference human proteome into phosphorylation likelihood sets based on available phosphorylation evidence in PhosphoSitePlus (PSP) and PeptideAtlas (PA).

| Phosphorylation likelihood set | Phosphorylation evidence per site | Ser count | Thr count | Tyr count |
| --- | --- | --- | --- | --- |
| High in PSP | 5+ pieces of evidence | 32306 | 8161 | 9763 |
| Medium in PSP | 2-4 pieces of evidence | 34154 | 12197 | 7228 |
| Low in PSP | 1 piece of evidence | 66777 | 35173 | 20191 |
| High in PA | 5+ observations at PTM score >0.95 which is $\geq 10\%$ of total observations | 26186 | 4204 | 1169 |
| Medium in PA | 5+ observations at PTM score >0.95 which is <10% of total observations OR 2-4 observations at PTM score >0.95 | 20517 | 5297 | 1460 |
| Low in PA | 1 observation at PTM score >0.95 | 12950 | 4895 | 1324 |
| Not phosphorylated | 0 observations at PTM score >0.19 AND 5+ observations at PTM score $\leq 0.19$ AND no evidence in PSP | 13892 | 8462 | 2184 |
| Other sites | At least 1 observation in PA but does not fall into any other PA categories AND no evidence in PSP | 60221 | 35000 | 10009 |

**Table 2.**
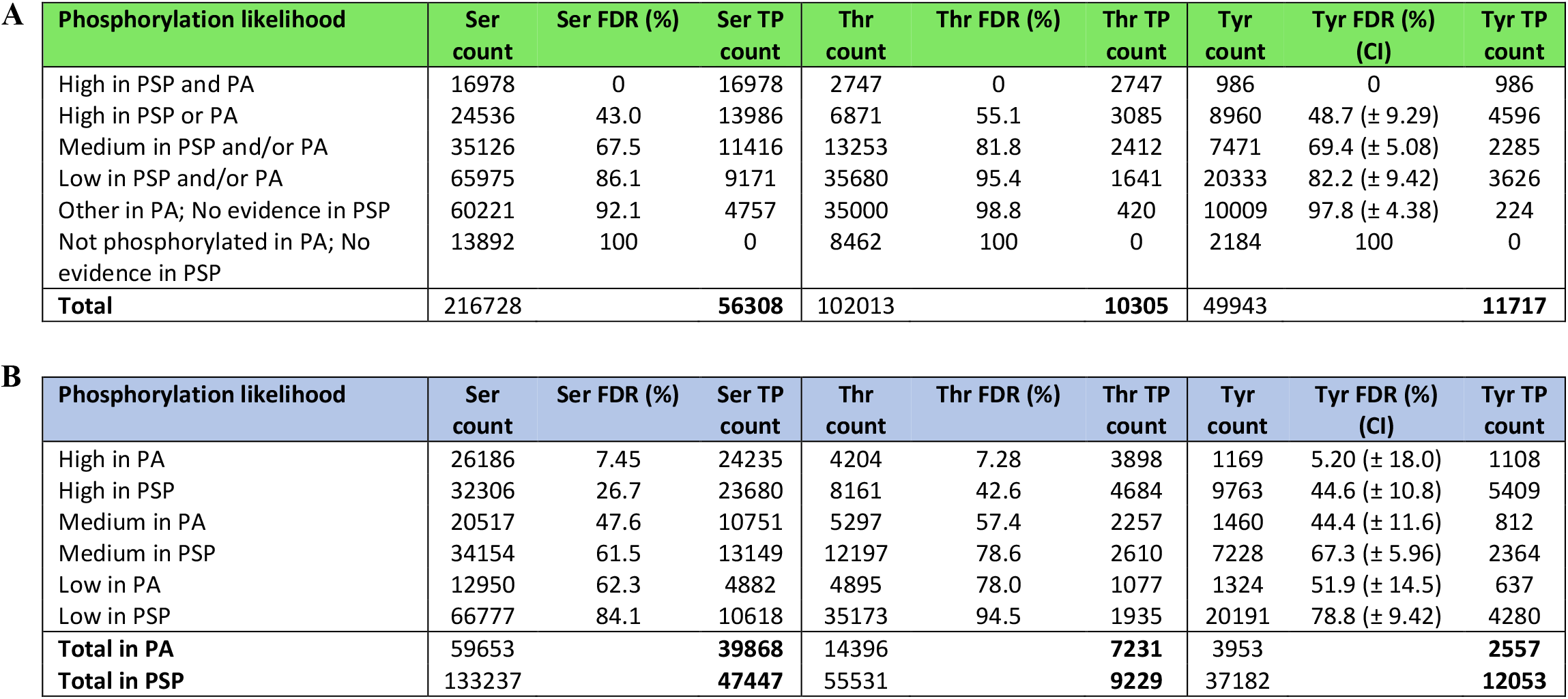
Counts of estimated true positive (TP) serine (Ser), threonine (Thr) and tyrosine (Tyr) phosphosites within sets of various phosphorylation likelihood based on **(A)** overall combined evidence and **(B)** individual positive identification evidence in PhosphoSitePlus (PSP) or PeptideAtlas (PA). Per each set, TP counts were derived from the FDR estimates within the set and the overall count of target amino acids in the set.

### Evolutionary conservation analysis

To determine the cross-species conservation of all Ser, Thr and Tyr sites in the reference human proteome (The UniProt, 2019) which have phosphorylation evidence in PSP and PA, human reference proteome (20,605 sequences, UniProt ID: UP000005640) and the proteomes of 100 eukaryotic species (50 mammals, 12 birds, 5 fish, 4 reptiles, 2 amphibians, 11 insects, 4 fungi, 7 plants and 5 protists; SI Table 3) were downloaded from UniProt (UniProt release 2019_10). Each sequence in the human proteome was used as a query in a BLASTp search (BLAST+ 2.10.0 version) (Altschul *et al*., 1990) against all 100 eukaryotic proteomes. The BLAST output (SI Table 4) was processed to extract a top matching significant orthologue (*E*-value of ≤ 0.00001) from each species for each human target. Human targets were then aligned with their matched orthologues using the MUSCLE algorithm (version 3.8.31) (Edgar, 2004) with default settings if all sequences to be aligned were <2,000 amino acids long. If any sequences to be aligned (either the human sequence or any of the orthologue sequences) were ≥2,000 amino acids long, 2 iterations of the algorithm were run using settings for large alignments (`-maxiters 2` option) (Edgar, 2004). From the alignments, percentage conservation scores were calculated for every Ser, Thr and Tyr site within each human target out of 100 (all eukaryotic proteomes) and out of the number of aligned orthologues. Conservation percentages were calculated considering any Ser/Thr substitutions in orthologues, whereby an orthologue was included in the count if, for example, a Thr in its sequence was aligned with a Ser in the target human sequence and vice versa. Conservation data was then cross-referenced with PSP/PA datasets to identify sites in the human proteome with phosphorylation evidence in PSP/PA and determine their conservation. To ensure consistency in terms of proteins and sites used, any human protein target for which it was not possible to calculate site conservation either due to the protein having no matches in BLAST (14 proteins), no significant matches in BLAST (236 proteins), no Ser/Thr/Tyr sites in its sequence (1 protein) or due to failed alignments (10 proteins), was excluded from any further analysis (SI Table 5). Any human targets labelled with the same UniProt identifier in the reference human proteome, PSP and PA, but which corresponded to different protein sequences across the datasets (73 proteins; SI Table 6) were also excluded. Conservation was assessed for the remaining targets (SI Material 1) by linear regression models with non-assumed intercept for simpler interpretation of slope between phosphosites and non-phosphosites. Average conservation of likely phosphosites (sites ranked “*High*” or “*Medium*” in PSP and/or PA) was plotted against average conservation of likely non-phosphosites (sites in “*Not phosphorylated*” and “*Other*” sets) within each target protein that had at least 3 likely phosphosites and 3 likely non-phosphosites. Conservation scores (%) were also compared across all sites within phosphorylation likelihood sets using box plots.

### Analysis of amino acids adjacent to phosphosites

Target protein sequences (20,271 sequences; SI Material 2) were processed to identify amino acids at the -1 and +1 proximal positions adjacent to every Ser, Thr and Tyr site. If a target sequence ended with a Ser, Thr or Tyr site then its +1 amino acid was marked as “*Not found*”. For each amino acid, its frequency at each proximal position was first normalised to 1,000 and then to its frequency in the pre-filtered human reference proteome (expected distribution). Proximal amino acid frequencies around target Ser, Thr and Tyr in “*High in PSP and PA*” set were compared to those in the “*Not phosphorylated*” set, and to the expected amino acid distribution. The comparisons were assessed by Fisher’s exact statistical test (Fisher, 1934) performed using scipy module in Python (Virtanen *et al*., 2020) with Bonferroni corrections to generate adjusted p-values. For each amino acid, any significant difference (Bonferroni corrected p-value <0.01) between the compared sets was used to estimate phosphosite false discovery rate across all phosphorylation likelihood sets. FDR estimates assumed that all sites in the highest phosphorylation likelihood set “*High in PSP and PA*” set were true positive phosphosite identifications, whereas all sites with the weakest phosphorylation confidence (either the “*Not phosphorylated*” or the “*Other*” set) were non-phosphosites. Therefore, the observed count of a certain proximal amino acid in the “*High in PSP and PA*” (nPos) corresponded to its expected count at 0% FDR, whereas its observed count in the “*Not phosphorylated*” or “*Other*” set (nNeg) corresponded to its expected count at 100% FDR. To estimate % FDR in any other phosphorylation likelihood set based on the observed count of the compared proximal amino acid in that set (nObs), we used the following equation:

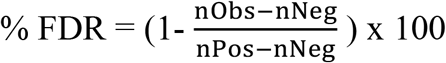

The equation has the effect of estimating what proportion of the observed count (nObs) is explained by assumed false positives (nNeg) and what proportion by true positives (nPos). For example, if amino acid X was found at +1 position next to 500 Ser sites in the highest phosphorylation confidence set (0% FDR set; nPos = 500) compared to 10 Ser sites in the “*Not phosphorylated*” set (100% FDR set; nNeg = 10), and next to 350 sites in the set of interest (nObs = 350), then pSer FDR within the set of interest would be 30.6%. This would suggest that 30.6% of sites in that set behave like false positive pSer in terms of X amino acid frequency at +1 position, whereas 69.4% of those sites behave like sites in the highest phosphorylation likelihood set (true pSer).

An average FDR with 95% confidence intervals (CI) was calculated per each likelihood set if multiple amino acids were significantly enriched at adjacent positions around a particular target phosphosite. Final FDR estimates were used to derive the total number of true positive (TP) phosphosite identifications across phosphorylation likelihood sets.

To compare FDR/TP estimates between individual PSP and PA sets, the method was replicated using alternative phosphorylation likelihood sets, where sites were categorised according to the highest amount of positive phosphorylation evidence from one database (at least 1 observation at PTM probability >0.95 in PA; at least 1 observation in PSP), without taking into account any evidence in the other. Phosphosite FDR estimates within “*High*” sets in each database were presented as a weighted average between FDR estimates in sites ranked “*High*” in that database only and sites ranked “*High*” in both PSP and PA. For example, the FDR in “*High in PA*” set was a weighted average of FDR estimates in “*High in both*” set and “*High in PA only*” set.

### Functional enrichment analysis

All protein sequences in the filtered reference human proteome (SI Material 2) were categorised into sets according to what their highest ranked Ser, Thr and Tyr site was in terms of phosphorylation evidence (“*High in PSP and PA*”, “*High in PSP or PA*”, “*Medium in PSP and/or PA*”, “*Low in PSP and/or PA*”, “*Other in PA*”, “*Not phosphorylated”* and “*No evidence in PSP or PA*”). Each protein set within Ser, Thr and Tyr datasets (SI Material 1) was analysed with DAVID (version 6.8) (Dennis *et al*., 2003) using all proteins in filtered proteome with any Ser, Thr or Tyr evidence in PSP or PA (16,296, 14,565 and 12,912 proteins respectively) as control background. Protein sets containing no reported evidence in PSP or PA were searched against a background of all proteins in the filtered reference proteome to determine any differences in their functional enrichment compared to proteins with PSP/PA evidence. Per each set searched, the top 10 (where possible) significant (Benjamini–Hochberg corrected p value <0.05) functional terms with the highest percentage of proteins mapped were identified, replacing any near synonymous terms with additional terms from outside the initial top 10. All target protein sets were also searched in UniProt (release 2020_04) to determine percentage of proteins mapped to UniProt keywords “*Phosphoprotein*”, “*Alternative splicing*”, “*Nucleus*”, “*Transcription*”, “*Acetylation*”, “*Membrane*”, “*Glycoprotein*”, “*Signal*” and “*Disulfide bond*”.

### Secondary structure analysis

Categorised Ser, Thr and Tyr sites in filtered reference human proteome were mapped to protein structures (beta strand, helix, turn and coiled coil) described for those proteins in UniProt (release 2020_04) (SI Material 1, SI Table 7). Any target proteins searched in UniProt which were marked as obsolete (15 proteins) or represented different sequences despite being labelled with the same identifier (25 proteins) were removed from the analysis and marked as “*NA*” (SI material 1). Normalised (to 1,000) counts of target amino acids within protein structures were assessed with Fisher’s exact statistical test (Fisher, 1934) using the scipy module in Python (Virtanen *et al*., 2020) to generate p-values and indicate any significant enrichment (p <0.05) between “*High in PSP and PA*” set and the “*Not phosphorylated*” set.

## Results and discussion

### Categorising all Ser, Thr and Tyr annotated phosphosites in the human proteome

We first ranked all Ser, Thr and Tyr phosphosites in PA and PSP in the filtered reference human proteome according to the amount of accumulated identification evidence (Fig. 1; Table 1; SI material 1). We found that the majority of Ser, Thr and Tyr sites (50.1%, 63.3% and 54.3% respectively) with phosphorylation evidence in PSP were placed into the “*Low*” phosphorylation likelihood set meaning that there was only 1 piece of evidence describing their positive identification (Fig. 1a). Furthermore, out of all analysed Ser, Thr and Tyr sites with at least 1 observation at PTM probability >0.95 in PA (suggesting a positive phosphosite identification), 21.7%, 34.0% and 33.5% respectively were placed in the “*Low*” set (Fig. 1b, Table 1), highlighting that a considerable amount of potential phosphosites only had 1 piece of positive identification evidence across both databases.

**Figure 1.**
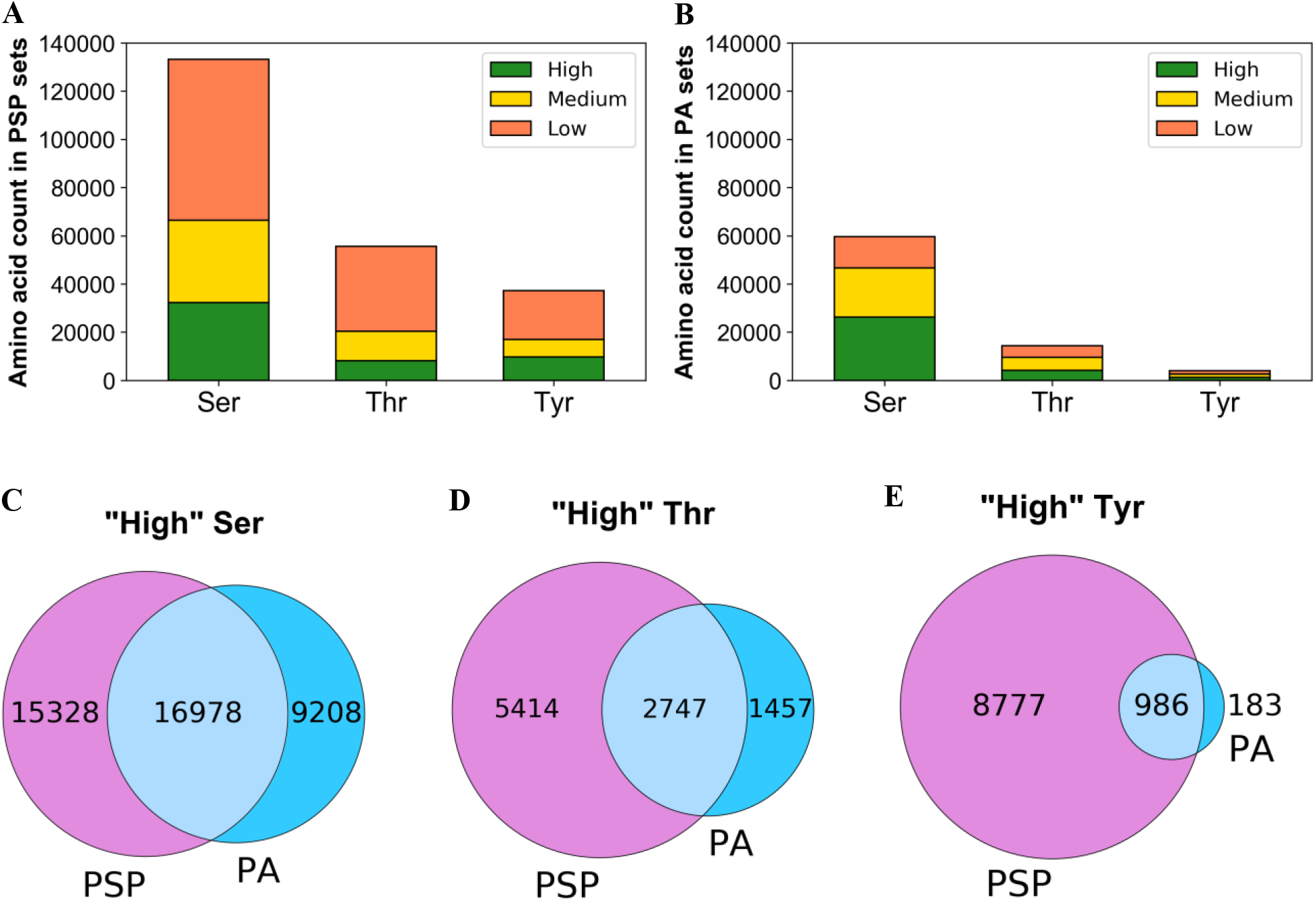
Distribution of serine (Ser), threonine (Thr) and tyrosine (Tyr) phosphosites from UniProt’s reference human proteome that have any positive identification evidence in **(A)** PhosphoSitePlus (PSP) or **(B)** PeptideAtlas (PA) based on established phosphorylation likelihood sets (see “Methods”). Venn diagrams provide the counts of **(C)** Ser, **(D)** Thr and **(E)** Tyr sites ranked “*High*” in PSP (left), PA (right) and both resources (overlap).

Interestingly, we found that in the human proteome there were more Tyr sites assigned to “*High*” set in PSP (5+ observations) than Thr sites (Fig. 1a, Table 1). Assuming that sites in the “*High*” set are likely to be real phosphosites, this finding contradicts the widely accepted notion that Thr phosphosites are more abundant than Tyr phosphosites (Hardman *et al*., 2019; Olsen *et al*., 2006), although this may vary depending on the active cell cycle phase or exposure to growth-factors (Caron *et al*., 2016). High prevalence of likely true Tyr phosphosites in the PSP dataset could have been a result of in-house studies which identified large numbers of pTyr sites using immunoaffinity strategies not suitable for pSer/pThr discovery (Rikova *et al*., 2007; Rush *et al*., 2005), and studies which have not been officially published (Hornbeck *et al*., 2015).

From PA, it is possible to identify sites for which covering phosphopeptides are observed but for which the modifications are only localised to other sites in the same peptides, thus providing strong evidence for likely non-phosphosites. Sets of potential Ser, Thr and Tyr non-phosphosites were therefore established based initially on evidence in PA (SI Table 8). Those sets were then cross-referenced with data in PSP to determine whether PSP contained any sites ranked as non-phosphosites in PA. Interestingly, we found that 2,489 Ser, 1,341 Thr and 891 Tyr sites assigned to the “*Not phosphorylated*” set in PA were found to have evidence in PSP (SI Table 9). In fact, out of those potential PA non-phosphosites, 146 Ser, 97 Thr and 293 Tyr sites were placed into “*High*” phosphorylation likelihood set according to PSP evidence (SI Table 9). This strongly indicated the presence of potential false positives in PSP and/or false negatives in PA. For example, sites Ser42 in protein P17066 (HSPA6) and Ser59 in Q8N488 (RYBP) had 8 and 6 phosphosite identification references in PSP respectively (mostly from in-house MS studies) but had no positive identification evidence in PA or any mention in other databases including UniProt (The UniProt, 2019) and neXtProt (Zahn-Zabal *et al*., 2020) (SI material 1). On the other hand, site Ser4 in P15927 (RPA2) had 33 phosphosite identification references in PSP and was also mentioned in UniProt’s and neXtProt’s annotations, but has never been positively localised in any of its 127 associated PSMs in PA (SI material 1). To eliminate potential false assignments when considering evidence in both PSP and PA, a site was only categorised as a non-phosphosite if it had no evidence in PSP in addition to having “negative” phosphorylation evidence in PA (Table 1). As a result, we established final negative control sets containing 13,892 Ser, 8,462 Thr and 2,184 Tyr sites. Similar adjustments were made to the “*Other*” PA set (sites in that set must have no evidence in PSP) which contained the majority of analysed PA sites (Table 1).

Having further cross-referenced sets of sites of various phosphorylation likelihood between PSP and PA (SI Table 9), we established a “gold standard” set of phosphosites, all of which had “*High*” phosphorylation likelihood according to both PSP and PA evidence. This set contained 16,978 Ser (Fig. 1c), 2,747 Thr (Fig. 1d) and 986 Tyr (Fig. 1e) highly likely true phosphosites. As for the general agreement between PSP and PA in terms of phosphorylation evidence, we found that 37.7% of Ser, 20.5% of Thr and 9.10% of Tyr sites with PSP evidence also had at least 1 observation at PTM probability >0.95 in PA (SI Table 9). This variation in sites observed between the two databases can be explained by the likely use of different methods for phosphosite detection and localisation between PA and the sources referenced in PSP, as well as due to a considerable presence of random false positives in both datasets before thresholding has been applied (see ‘Introduction’).

### Evolutionary conservation analysis

Phosphoproteomes from all species are constantly evolving, although many ancient phosphosites are conserved across species and taxa, where they are likely to be functionally relevant (Boekhorst *et al*., 2008; Malik, Nigg and Körner, 2008; Studer *et al*., 2016). In our analysis we determined the conservation of all potential Ser, Thr and Tyr phosphosites and non-phosphosites in UniProt’s filtered reference human proteome across 100 eukaryotic species, weighed towards vertebrates, but also including examples of insects, plants and unicellular eukaryotes (SI Table 3). In our first analysis, we explored the mean conservation of phosphosites and non-phosphosites per protein (at least three of each per protein) and performed a correlation analysis across all proteins (Fig. 2). We fitted linear regression models through the origin, under the theory that proteins unique to humans would have zero conservation for both phosphosites and non-phosphosites. We found great variation between the conservation of both site types, ranging from near zero to 100%, which was mostly dependent on the overall conservation of the protein sequence. However, based on the generated linear regression models, we concluded that on average, Ser, Thr and Tyr phosphosites (“*High*” or “*Medium*” in PSP and/or PA) were around 4.6%, 5.4%, and 2.0% respectively more conserved across all 100 eukaryotes than corresponding likely non-phosphosites (sites in “*Not phosphorylated*” and “*Other*” sets) within analysed proteins when allowing Ser/Thr substitutions towards the conservation score (Fig. 2). Similar results were obtained when assessing phosphosite conservation only across found orthologues for each protein (SI Fig. 1). The results (Fig. 2, SI Fig. 1) provide additional evidence that phosphosites are generally more conserved that non-phosphosites (Boekhorst *et al*., 2008; Chen, Chen and Li, 2010; Malik, Nigg and Körner, 2008). The difference in conservation is thus subtle and variable, but statistically robust. Furthermore, in our analysed sets of proteins which had at least 3 likely phosphosites and 3 likely non-phosphosites, we found 104, 88 and 19 proteins where the conservation of Ser, Thr and Tyr likely phosphosites respectively was at least 20% higher than conservation of likely non-phosphosites (SI Table 10).

**Figure 2.**
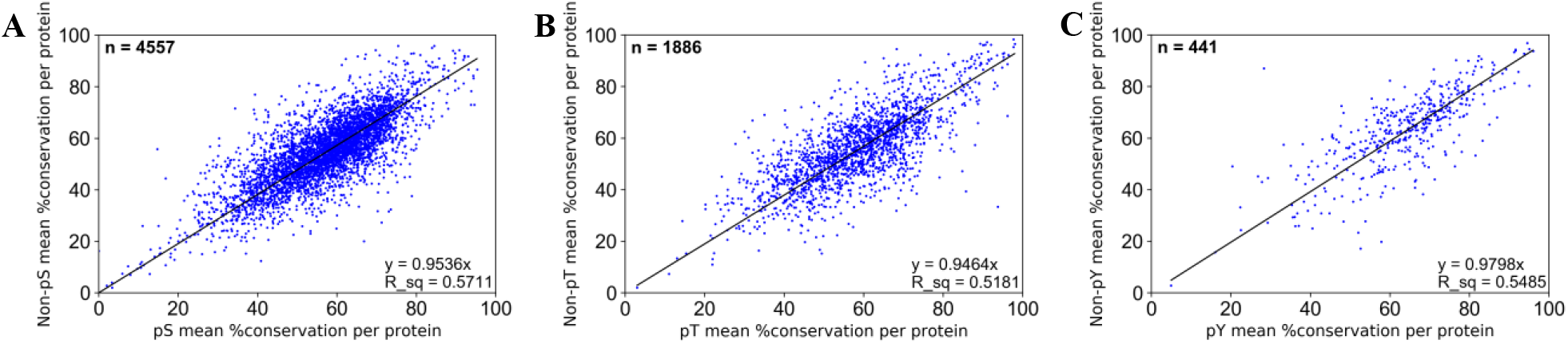
Mean % conservation across 100 eukaryotic species of likely **(A)** Ser, **(B)** Thr, **(C)** Tyr phosphosites and corresponding likely non-phosphosites within each target protein (n = number of proteins analysed). The regression coefficient (R^2^) is given by ‘R_sq’.

In our next analysis, we compared the conservation of all sites split by phosphorylation likelihood sets (Fig. 3) and revealed that sites in the highest phosphorylation likelihood set (“*High in both PSP and PA*”) had the highest average conservation across all 100 eukaryotic proteomes considering Ser/Thr substitutions (average conservation of 58.4%, 58.6% and 69.4% across 16,978 Ser, 2,747 Thr and 986 Tyr sites respectively) (Fig. 3, SI Table 11). In comparison, the sites in “*Low in PSP and/or PA*” set had slightly lower average conservation scores of 54.3%, 55.4% and 64.0% in 35126 Ser, 13253 Thr and 7471 Tyr sites respectively (Fig. 3, SI Table 11). Assuming that high conservation is a property of true phosphosites, the results (Fig. 3) show that this property was observed more frequently in higher phosphorylation likelihood sets compared to lower ones suggesting higher potential phosphosite FDR in sets with weaker phosphorylation evidence.

**Figure 3.**
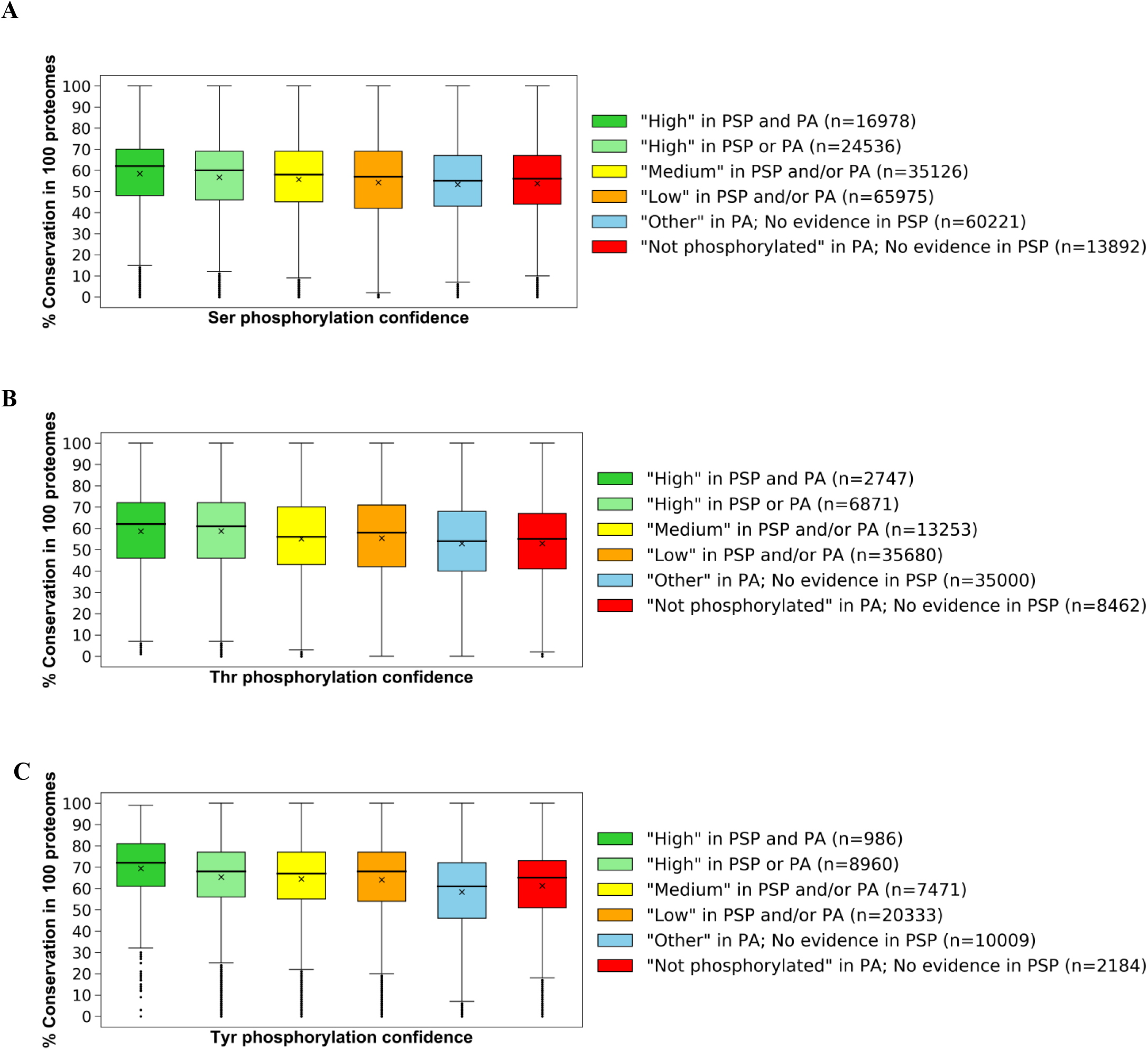
Box plots of conservation percentages (%) across 100 eukaryotic species of human **(A)** Ser, **(B)** Thr and **(C)** Tyr sites categorised across established phosphorylation confidence sets based on PSP and PA evidence. Within each box, a horizontal line represents median % conservation, an (x) symbol represents mean % conservation per group. Each box extends from the 25th to the 75th percentile of each set’s distribution of conservation % values. Vertical lines extending from the boxes correspond to adjacent values. Dots (•) represent outlier values.

There were numerous cases in our analysis of likely non-phosphosites and sites with “*Low*” phosphorylation likelihood where amino acid conservation was also high compared to likely phosphosites (SI material 1), indicative of a conserved function for these amino acids in, for example, catalysis or a biomolecular interaction that is unrelated to phosphorylation. Furthermore, we found 64, 30 and 6 proteins in which the average conservation across 100 eukaryotes of Ser, Thr and Tyr likely non-phosphosites respectively was at least 20% higher than the conservation of corresponding likely phosphosites (SI Table 10). It is possible that the predicted phosphosites within those proteins were either false positives or were non-functional true phosphosites, explaining the comparative weaker selective pressure. In fact, previous reports estimated that as many as 65% of known phosphosites may be non-functional due to limited kinase specificity and therefore have similar evolution rates compared to non-phosphosites which would explain the trends in our analysis (Landry, Levy and Michnick, 2009; Lienhard, 2008). It is also possible that some proteins could have been formed by recent gene fusion events leading to regions containing phosphorylation sites only found in a few closer related orthologues (low conservation), with other protein domains being more highly conserved.

### Analysis of amino acids adjacent to phosphosites

Amino acids directly adjacent to known phosphorylation sites are often involved in optimising substrate capture for subsequent phosphotransfer by the kinase enzymatic machinery (Byrne *et al*., 2020; Hutti *et al*., 2004; Kettenbach *et al*., 2012). In our analysis we identified the frequency of -1 and +1 amino acids relative to a possible phosphosite and compared it across different sets of sites ranked by the relative strength of phosphorylation evidence (Table 1). We found a strong enrichment of proline (Pro) at the +1 position next to Ser and Thr sites in the reference human proteome that were placed in the set with the most phosphorylation evidence (“*High” in PSP and PA*”) (Fig. 4a & 4b, SI Table 12). In fact, Pro was observed at the +1 position next to 44.3% and 74.9% of all Ser and Thr sites respectively in that set (SI Table 12). The enrichment of Pro at +1 position around those sites was significant (adj. p value < 0.01) in relation to the normalised distribution of Pro in the human proteome, where it is, in fact, only the sixth most observed amino acid (SI Table 12). The normalised number of +1 observations of Pro around Ser and Thr sites in the highest phosphorylation likelihood set was also significantly (adj. p value <0.01) higher than around Ser and Thr sites in the “*Not phosphorylated*” set (Fig. 4a & 4b), where only 2.68% of Ser and 5.67% of Thr sites had Pro at +1 position (SI Table 12). Therefore, the enrichment of Pro around highly likely Ser and Thr phosphosites suggests that this feature, amongst others, can be used as a differentiating characteristic for phosphosites compared to non-phosphosites. Furthermore, multiple previous reports specifically highlight the importance of Pro in the mechanism of phosphorylation for families of kinases such as the cyclin-dependent kinases, Mitogen Activated Protein Kinases and, more recently, the centrosomal kinase PLK4 (Byrne *et al*., 2020; Hall and Vulliet, 1991; Johnson *et al*., 1998; Keshwani *et al*., 2015; Lu, Liou and Zhou, 2002; Pietrangelo and Ridgway, 2019; Songyang *et al*., 1994). Consequently, there is a high prevalence of Pro in numerous phosphorylation motif sequences as part of Ser/Thr-Pro combinations (Amanchy *et al*., 2007; Sugiyama, Imamura and Ishihama, 2019). As a result, we compared the frequency of Pro across the sites within different phosphorylation likelihood sets and used the comparison to estimate phosphosite false discovery rate across those sets. We found that in both Ser and Thr datasets, the number of Pro observations at +1 position next to target sites decreased with the amount of phosphorylation evidence, suggesting higher phosphosite FDR in Ser/Thr sets with weaker phosphorylation evidence (Fig. 4a & 4b). Using the counts of Pro at +1 position around Ser sites of various phosphorylation likelihood (SI Table 12) and working under the assumption of FDR = 0% in set 1 “*High” in PSP and PA*”, we estimated Ser phosphosite FDR = 43.0% in set 2 “*High” in PSP or PA”*; FDR = 67.5% in set 3 “*Medium in PSP and/or PA*”; FDR = 86.1% in set 4 “*Low in PSP and/or PA*” and FDR = 92.1% in the “*Other*” set. Using the same method for the Thr data, we estimated Thr phosphosite FDR = 55.1% in set 2; FDR = 81.8% in set 3; FDR = 95.4% in set 4 and FDR = 98.8% in the “*Other*” set. Our FDR estimates clearly highlight that the majority of Ser and Thr sites with just 1 piece of phosphosite identification evidence are likely false positive identifications and users of these databases can reasonably assume that if a site does not have multiple levels of evidence, then it is unlikely to represent a true phosphorylation site.

**Figure 4.**
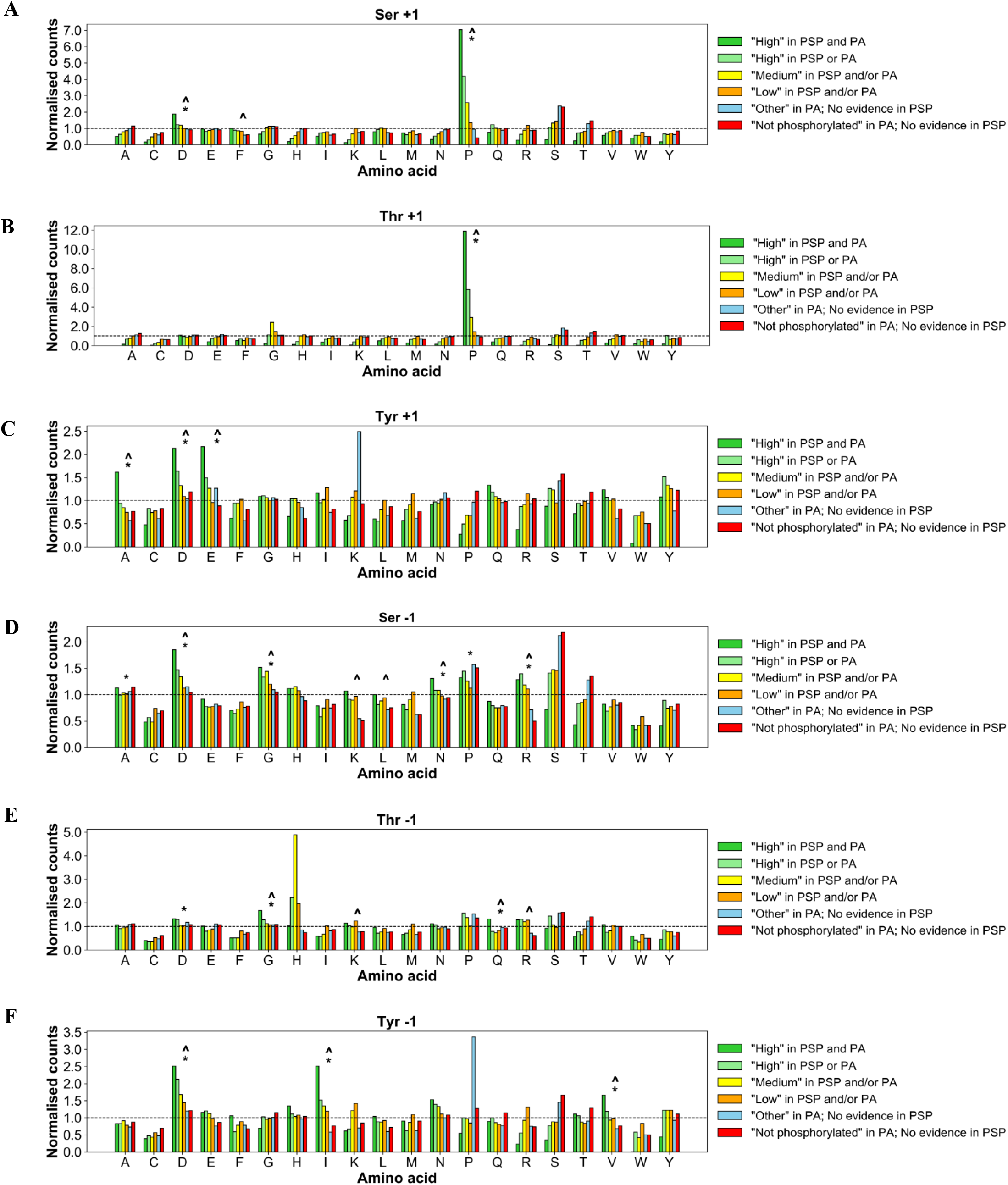
Counts of proximal amino acids positioned at **(A)** +1 around Ser; **(B)** +1 around Thr; **(C)** +1 around Tyr; **(D)** -1 around Ser; **(E)** -1 around Thr; **(F)** -1 around Tyr sites of various phosphorylation likelihood based on evidence in PSP and PA, normalised to observed distribution of those amino acids in human proteome (represented by dotted baseline set at 1). Significant (Bonferroni corrected p value <0.01) enrichment of proximal amino acids in the “*High” in PSP and PA*” set is highlighted by the caret symbol (**^**) when compared against the “*Not phosphorylated*” set, and an asterisk symbol (*) when compared to the expected amino acid distribution.

We also found a significant enrichment of aspartic acid (Asp or D) at +1 position next to Ser sites in the highest phosphorylation likelihood set (Fig. 4a). To explain this, we linked the sequences containing those sites to Casein kinase II phosphorylation motifs which commonly feature Ser-Asp combinations (Amanchy *et al*., 2007; Songyang *et al*., 1996). At the -1 positions around target Ser, we found significant enrichment (adj. p value <0.01) of Asp, glycine (Gly or G), asparagine (Asn or N) and arginine (Arg or R) in the highest phosphorylation likelihood set compared to “*Not phosphorylated*” set (Fig. 4d). It is possible that the observed enrichment was due to the presence of those amino acids within substrate motifs of Casein Kinase II, CDK5, PKC, PKA and MEKK (Amanchy *et al*., 2007), suggesting high prevalence of potential true Ser phosphosites. Similar conclusions were made for the enrichment of Gly at -1 around Thr sites in the highest phosphorylation likelihood set (Fig. 4e) which was linked to possible Gly-Thr combinations within PKA, ERK1 and ERK2 kinase substrate motifs (Amanchy *et al*., 2007). The enrichment of highlighted amino acids at the -1 position around Ser/Thr sites in the highest phosphorylation likelihood set (Fig. 4d & 4e) was not as strong as the enrichment of Pro at +1 position (Fig. 4a & 4b), and not as well cited as Pro in relation to phosphorylation (Hall and Vulliet, 1991; Keshwani *et al*., 2015; Lu, Liou and Zhou, 2002; Pietrangelo and Ridgway, 2019). As a result, we only relied on Pro enrichment at +1 positions to maximise the accuracy of FDR estimations within the sets of Ser and Thr sites.

In our analysis of proximal sites around target Tyr, we found a significant enrichment (adj. p value <0.01) of alanine (Ala or A), glutamic acid (Glu or E) and Asp at +1 positions, in addition to enriched isoleucine (Ile or I), valine (Val or V) and Asp at -1 positions in “*High in PSP and PA*” set compared to “*Not phosphorylated*” set (Fig. 4c & 4f). We were able to link the enrichment of those proximal sites to their possible involvement in various phosphorylation motifs including EGFR and Abl kinase substrate motifs; PTP1B and PTPRJ phosphatase substrate motifs, and multiple SH2 domain binding motifs (Amanchy *et al*., 2007; Wälchli *et al*., 2004), therefore indicating higher frequency of true Tyr phosphosites in the highest confidence set compared to other sets. Since there was not a single proximal amino acid strongly enriched as, for example, Pro at +1 in Ser/Thr analysis, we used the frequencies of all six enriched proximal amino acids around target Tyr in “*High in PSP and PA*” (Fig. 4c & 4f, SI Table 12) to estimate average Tyr phosphosite FDR with 95% confidence intervals. We estimated FDR = 48.7% (CI ± 9.29%) in set 2 “*High in PSP or PA*”, FDR = 69.4% (CI ± 5.08%) in set 3 “*Medium in PSP and/or PA*”, FDR = 82.2% (CI ± 9.42%) in set 4 “*Low in PSP and/or PA*”, and FDR = 97.8% (CI ± 4.38%) in the “*Other*” set (Table 2a; SI Table 13).

Based on our Ser, Thr and Tyr phosphosite FDR estimates, we predicted that there were 56,308 Ser, 10,305 Thr and 11,717 Tyr true positive (TP) phosphosite identifications in the human proteome that were supported by evidence in PSP and/or PA (Table 2a). Furthermore, the results suggested that 91,064 Ser, 48,666 Thr and 26,257 Tyr sites with positive phosphorylation evidence in PSP and/or PA (sites in “*High*”, “*Medium*”, “*Low*” sets) were false positives (Table 2a). Interestingly, the estimated count of Tyr TPs was higher than the count of Thr TPs which goes against the general understanding of threonine phosphorylation being more prevalent than tyrosine (Olsen *et al*., 2006). One explanation for this could be that initially there were more Tyr sites with strong phosphorylation evidence than Thr sites, particularly in PSP (Fig. 1a, Table 2a). Furthermore, it is possible that some sites in the “*Not phosphorylated*” and “*Other*” sets were false negatives. In fact, we found 6,626 Thr sites which had no positive phosphorylation evidence in PSP or PA but were mentioned as phosphosites in UniProt and/or neXtProt (SI material 1), suggesting possible underestimation of true positive Thr phosphosites in our analysis.

Using the same method, the analysis of Pro frequency at +1 position adjacent to target Ser/Thr sites, and the frequency of enriched Ala/Glu at +1 and Asp/Ile/Val at -1 around target Tyr compared phosphosite FDR between PSP and PA sets by considering positive phosphorylation evidence (“*High*”, “*Medium*” or “*Low*” sets) in one database without taking into account any evidence in the other (SI Fig. 2, SI Table 14). The analysis revealed a generally lower FDR per each set in PA compared to the respective set in PSP, suggesting that a higher proportion of sites in the human proteome assigned to a certain phosphorylation likelihood set based on PA evidence only are true phosphosites compared to sites assigned to the same set based on only the PSP evidence (Table 2b).

### Functional enrichment analysis

In our analysis we categorised all 20,271 proteins in the filtered human reference proteome (SI Material 1) according to what their highest ranked Ser, Thr and Tyr site was based on phosphorylation likelihood sets in Table 1. The resulting sets (SI Table 15) were analysed in DAVID (Dennis *et al*., 2003) to compare functional enrichment patterns between phosphorylation likelihood sets. First, we found that across all datasets (Ser, Thr and Tyr) the protein sets containing sites ranked “*High in both PSP and PA*” were associated with the most significant (Benjamini–Hochberg adj. p value <0.05) functional groups (SI Fig. 3) suggesting their functional coherence i.e., sharing mappings to keywords, ontology terms or pathways. Interestingly, proteins with sites from “*Low in PSP and/or PA*” set as their highest ranked site and proteins which did not have any evidence phosphorylation evidence (“*No evidence in PSP or PA*” set) were also enriched for numerous functional categories suggesting that they too share some functional properties (SI Fig. 3). Proteins containing sites from the “*Not phosphorylated*” set as their highest ranked Ser/Thr/Tyr site were enriched for 1 significant functional group in the case of Tyr dataset and no functional groups in the case of Ser/Thr datasets, which was likely due to small protein sample size in those sets.

To investigate this further, we compared the top 10 enriched functional groups between the protein sets and found that proteins containing Ser, Thr and Tyr sites with most phosphorylation evidence (“*High in PSP and PA*” set) were significantly enriched for categories and terms associated with phosphorylation such as “*Phosphoprotein*”, “*Transcription*”, “*Nucleus*” and “*Alternative splicing*” (Fig. 5) suggesting that those proteins were true phosphoproteins. There is a risk of generating circular evidence here, as the enriched term “*Phosphoprotein*” is a UniProt keyword, and will have been annotated based on literature evidence, potentially shared with PSP. UniProt does not yet load phosphorylation evidence from high-throughput data sets, and so classifications of phosphoproteins are generally independent of evidence used in PA. Other enriched keywords have also likely been determined based on independent evidence, and thus we believe are unbiased observations of our sets. Overall, 92.3%, 93.9% and 88.2% of proteins containing Ser, Thr and Tyr sites of the highest phosphorylation likelihood respectively were enriched for the term “*Phosphoprotein*”, which, as per description in UniProt, is a term assigned to a “*protein which is post-translationally modified by the attachment of either a single phosphate group, or of a complex molecule, such as 5’-phospho-DNA, through a phosphate group*” (The UniProt, 2019). Furthermore, those proteins were enriched for “*Acetylation*” which in some cases might indicate phosphorylation since crosstalk between acetylation and phosphorylation has been frequently reported (Espinos *et al*., 1999; Habibian and Ferguson, 2018). In comparison, proteins that only had sites from “*Low in PSP and/or PA*” set as their highest ranked Ser, Thr and Tyr sites (i.e., proteins which did not have sites with strong phosphorylation evidence) were not enriched for clear phosphorylation-associated terms and were instead enriched for categories such as “*Glycoprotein*”, “*Signal*” and “*Disulfide bond*” and “*Membrane*” (Fig. 5), suggesting that the majority of those proteins were likely non-phosphoproteins and their associated phosphosites with weak evidence were therefore likely false positives. Assuming that sites with no phosphorylation evidence in PSP or PA are likely non-phosphosites (although it is possible that phosphorylation has not been investigated or localised yet), potential high FDR in the “*Low in PSP and/or PA*” set was further supported by proteins with no phosphorylation evidence being enriched for similar functional groups (Fig. 5). In fact, we observed a clear general decrease in percentage of proteins enriched for phosphorylation-associated functional groups (where a set was enriched for at least 10 functional groups) going across our established sets suggesting an increase in phosphosite FDR across the sets (SI Fig. 4).

**Figure 5.**
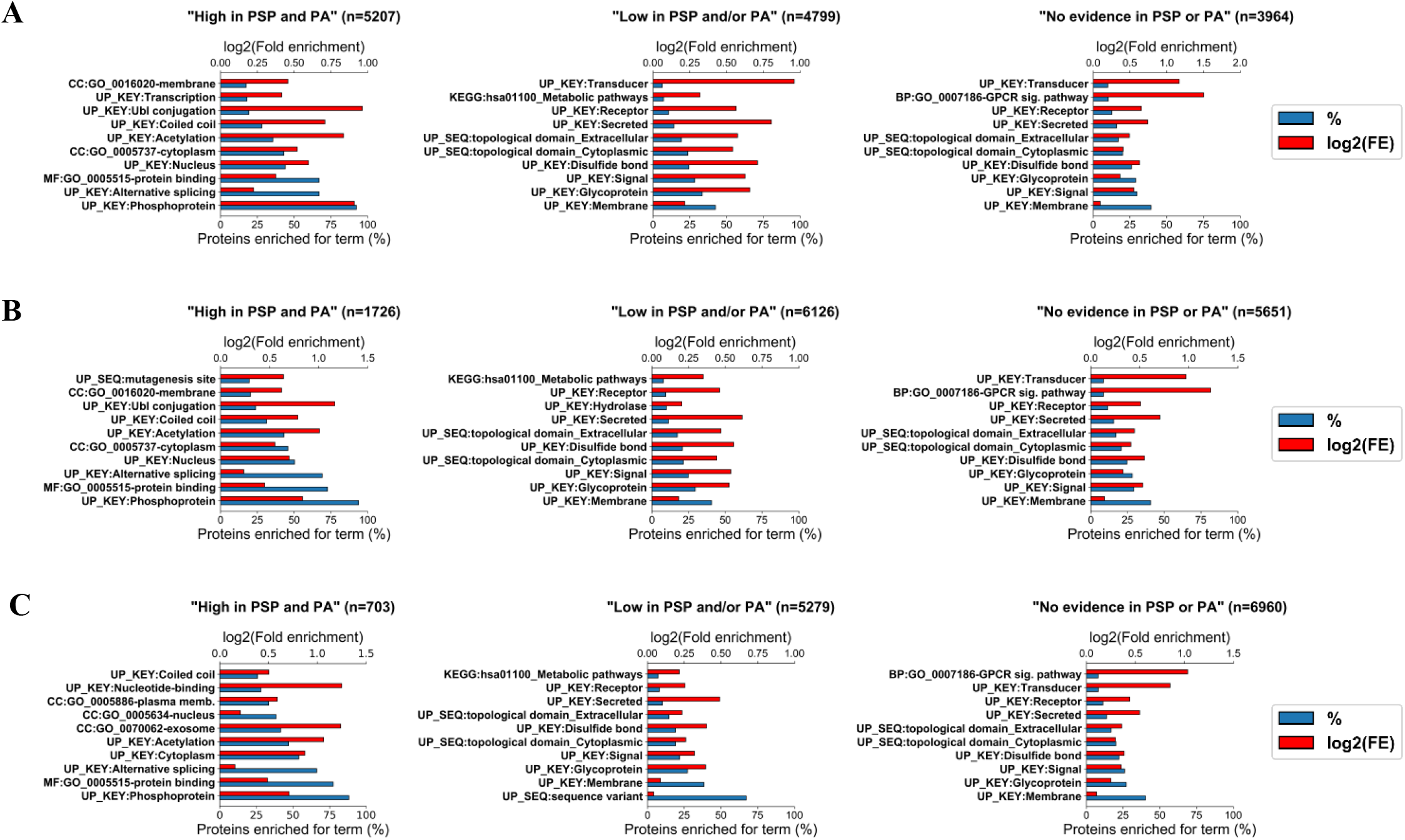
Top 10 functional categories for which protein sets containing various highest ranked **(A)** Ser, **(B)** Thr, **(C)** Tyr sites based on the amount of available phosphorylation evidence (“*High in PSP and PA*”, “*Low in PSP and/or PA*”, “*No evidence in PSP or PA*”) were significantly enriched in DAVID (Benjamini–Hochberg corrected p value <0.05). For each protein set, the % of proteins enriched for a particular functional category is given as well as the log2(fold enrichment) for that set. The number of proteins in each set is presented by n.

Our investigation of UniProt terms linked to protein sets revealed that the enrichment for term “*Phosphoprotein*” and other terms likely to be associated with phosphorylation (“*Alternative splicing*”, “*Nucleus*”, “*Acetylation*”, “*Transcription*”) generally decreased across confidence sets, which suggested higher FDR in sets with fewer phosphorylation evidence (Fig. 6; SI Table 16). For example, only 13.0%, 31.7% and 36.3% of all proteins, which had Ser, Thr and Tyr sites respectively from “*Low in PSP and/or PA*” phosphorylation likelihood set as their most confident site, were marked as phosphoproteins in UniProt (Fig. 6, SI Table 16), suggesting that most proteins in those sets were not phosphoproteins and further highlighting that the associated sites with 1 piece of phosphorylation evidence are likely false positive identifications.

**Figure 6.**
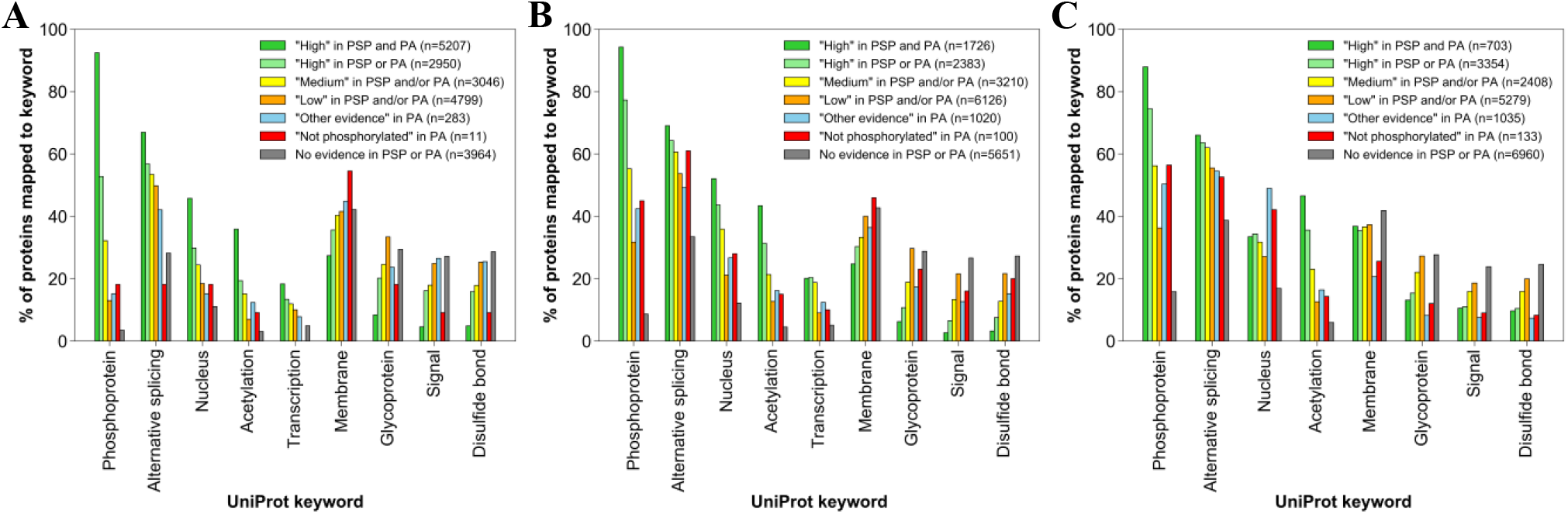
The percentage of proteins within sets containing **(A)** Ser, **(B)** Thr, **(C)** Tyr sites of various phosphorylation likelihood as their highest ranked site, annotated with specific UniProt keywords. The number of proteins in each set is presented by n.

### Secondary structure analysis

We also investigated whether Ser, Thr and Tyr sites with strong phosphorylation evidence were located more frequently within specific protein secondary structures, when compared to sites with less evidence. For example, previous analysis of thousands of phosphosites from multiple species identified hotspots within domain families of proteins, particularly near domain interfaces and adjacent to catalytic residues, where they presumably regulate enzymatic output (Beltrao *et al*., 2012; Strumillo *et al*., 2019). We found that significantly more (Fisher’s test p value <0.05) Ser, Thr and Tyr sites with the strongest phosphorylation evidence (“*High” in PSP and PA*” set) were localised within coiled coils compared to sites in the “*Not phosphorylated*” set (Fig. 7). This might readily be explained by coiled coils being frequently found in transcription factors, the activity or subcellular location of which is often dependent on phosphorylation (Barbara *et al*., 2007; Baxevanis and Vinson, 1993; Pogenberg *et al*., 2020). Therefore, the results in Fig. 7 further indicated that there were more potential true Ser, Thr and Tyr phosphosites in “*High” in PSP and PA*” set compared to “*Not phosphorylated*” set. In terms of other analysed protein structures (beta strand, turn, alpha helix), there was no significant enrichment of sites from the highest phosphorylation confidence set within those structures when compared to the “*Not phosphorylated*” set (Fig. 7). In fact, our current reading of the literature suggests that it is still unclear whether phosphorylation sites are found on average to be localised more or less frequently within beta strands, turns or alpha helices, though clear evidence for localisation of PTMs at functionally important loci in proteins has been presented (Gu and Wang, 2012).

**Figure 7.**
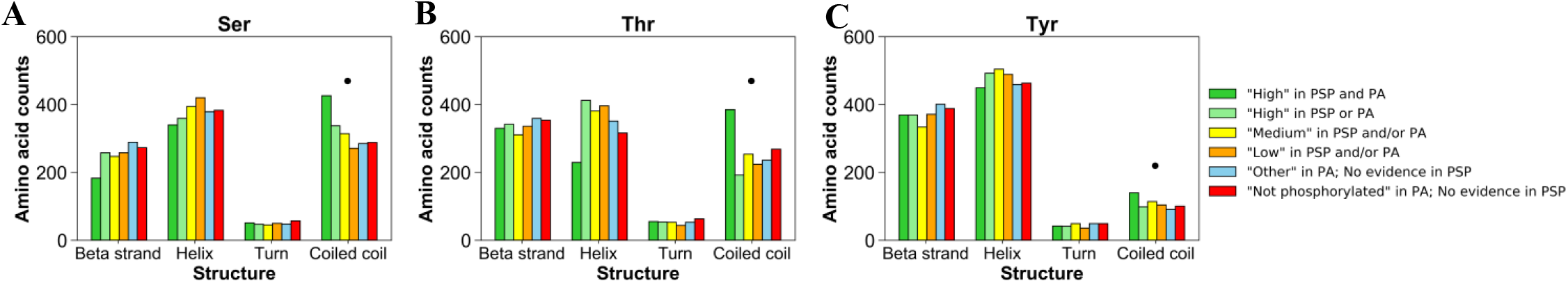
Normalised counts of **(A)** Ser; **(B)** Thr; **(C)** Tyr amino acids of various phosphorylation likelihood based on evidence in PSP and PA which are found within protein structures (beta strand, alpha helix, turn, coiled coil). Significant (Fisher’s test p value <0.05) enrichment of amino acids from “*High” in PSP and PA*” set within protein structures is highlighted by the dot symbol (•) when compared against the “*Not phosphorylated*” set.

## Conclusion

In our analysis we ranked all potential Ser, Thr and Tyr phosphosites in UniProt’s reference human proteome according to how much quantitative and qualitative phosphorylation evidence they were assigned in PSP and PA databases. Having analysed the sites and the proteins containing them in terms of conservation, proximal site patterns, functional enrichment and structural properties, we found that Ser, Thr and Tyr sites with weak phosphosite identification evidence, particularly sites that were only observed once, were likely to be false positive identifications. This finding was further confirmed by FDR estimations across the established phosphorylation likelihood sets which revealed phosphosite FDR of 86.1%, 95.4% and 82.2% in sets of Ser, Thr and Tyr sites respectively where only 1 piece of identification evidence was present. Since there is a considerable presence of such sites in PSP and PA datasets, our results implied high FDR in both those datasets, although PSP was predicted to have a generally higher proportion of false positive phosphosites compared to PA. This is potentially a cause for concern since many potential false positives are presented to scientists as true phosphosites, without clear presentation of the likelihood of such claims. Nevertheless, using our FDR estimates we predicted that there are 56308 Ser, 10305 Thr and 11717 Tyr true positive phosphosites in the human proteome that are supported by evidence in PSP and/or PA. These estimated counts are lower than other published estimates (Hornbeck *et al*., 2015; Ochoa *et al*., 2020; Safaei *et al*., 2011) particularly for Ser/Thr sites, presumably due to the previous inclusion of false positives and subsequent overestimation of the number of true phosphosites. We conclude that researchers must be aware of the potential for false positive sites in both public and self-generated databases and should always evaluate the evidence behind the phosphosites used in their research. We have provided here a methodological framework for estimating global FDR in large-scale phosphorylation data sets, which does not rely on native scores from search engines or site localisation software. Methods for estimating global FDR in meta-analyses of phosphosites are not yet robust, and thus we would recommend that other groups similarly profile orthogonal properties of ranked sets, as we have done here, to estimate the true and false phosphosite proportions in their data.

## Supporting information

SI_Material1_All_Data

SI_material2_Target_Protein_Sequences

SI_Fig1_conservation_in_orthologues_only

SI_Fig2_prox_site_plots_PSP_PA_separate

SI_Fig3_significant_group_count

SI_Fig4_all_sets_in_functional_analysis

SI_Tab1_PA_filtered_build

SI_Tab2_PSP_filtered_build

SI_Tab3_proteomes_table

SI_Tab4_blast_result_file

SI_Tab5_targets_not_analysed

SI_Tab6_STY_targets_with_sequences_not_matched

SI_Tab7_Secondary_structures_in_target_proteins

SI_Tab8_STY_counts_across_initial_PA_categories

SI_Tab9_cross_referencing_PSP_PA_categories

SI_Tab10_conservation_stats_within_proteins

SI_Tab11_conservation_stats_per_likelihood_set

SI_Tab12_proximal_site_data

SI_Tab13_FDR_calculations

SI_Tab14_proximal_site_data_PSP_PA_separate

SI_Tab15_ranked_proteins_for_functional_analysis

SI_Tab16_Uniprot_terms_analysis

