## Supplementary material for "Profiling the Human Phosphoproteome to Estimate the True Extent of Protein Phosphorylation": SI_Fig1_conservation_in_orthologues_only

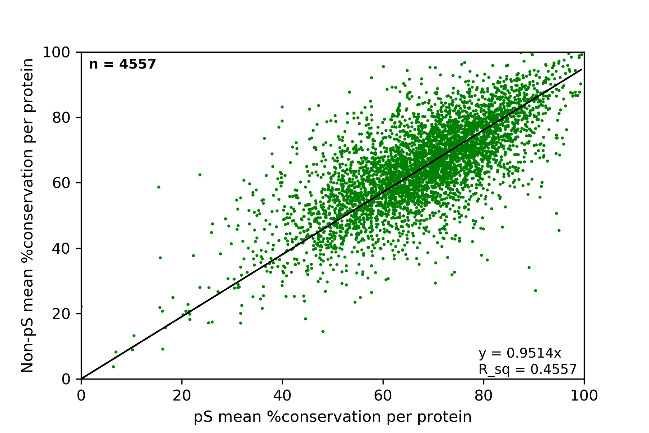

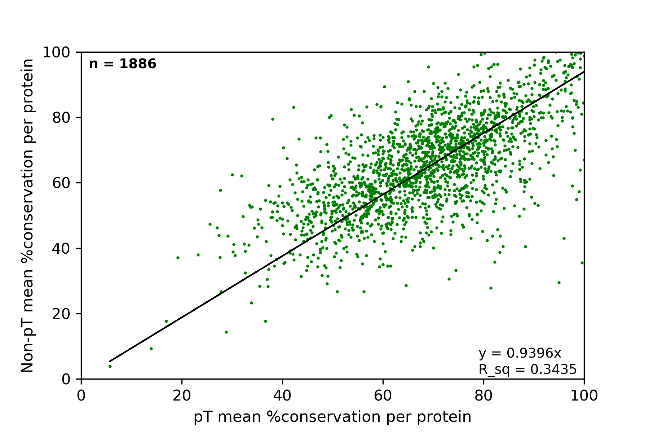

**B**

**A**

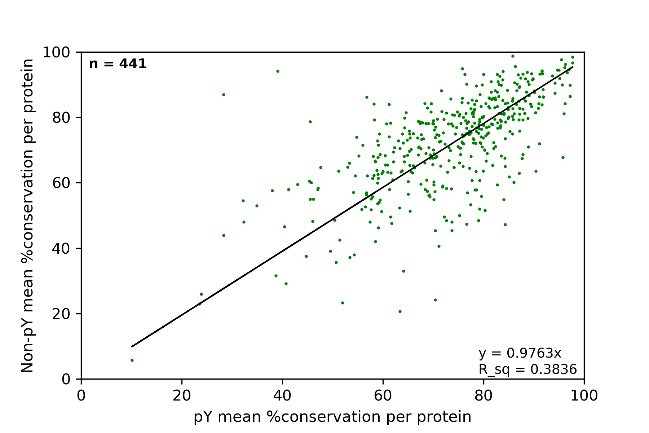

**C**

**SI Figure 1.** Mean % conservation across found orthologues of likely **(A)** Ser, (**B)** Thr, (**C)** Tyr phosphosites and corresponding likely non-phosphosites per each human protein in the analysed set (n). The R^2^ coefficient is given by ‘R_sq’.
