## Supplementary material for "Profiling the Human Phosphoproteome to Estimate the True Extent of Protein Phosphorylation": SI_Fig2_prox_site_plots_PSP_PA_separate

**A**

**B**

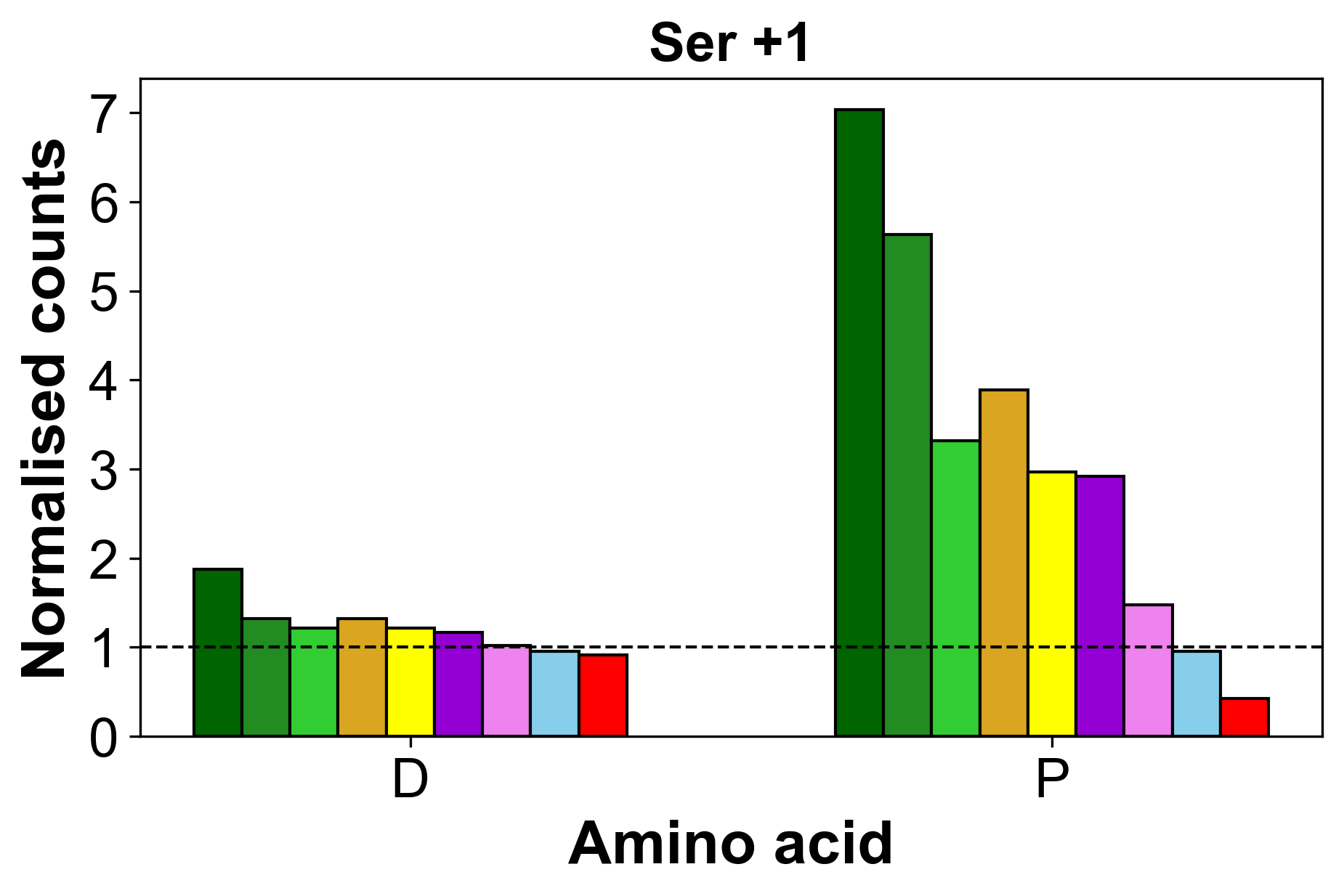

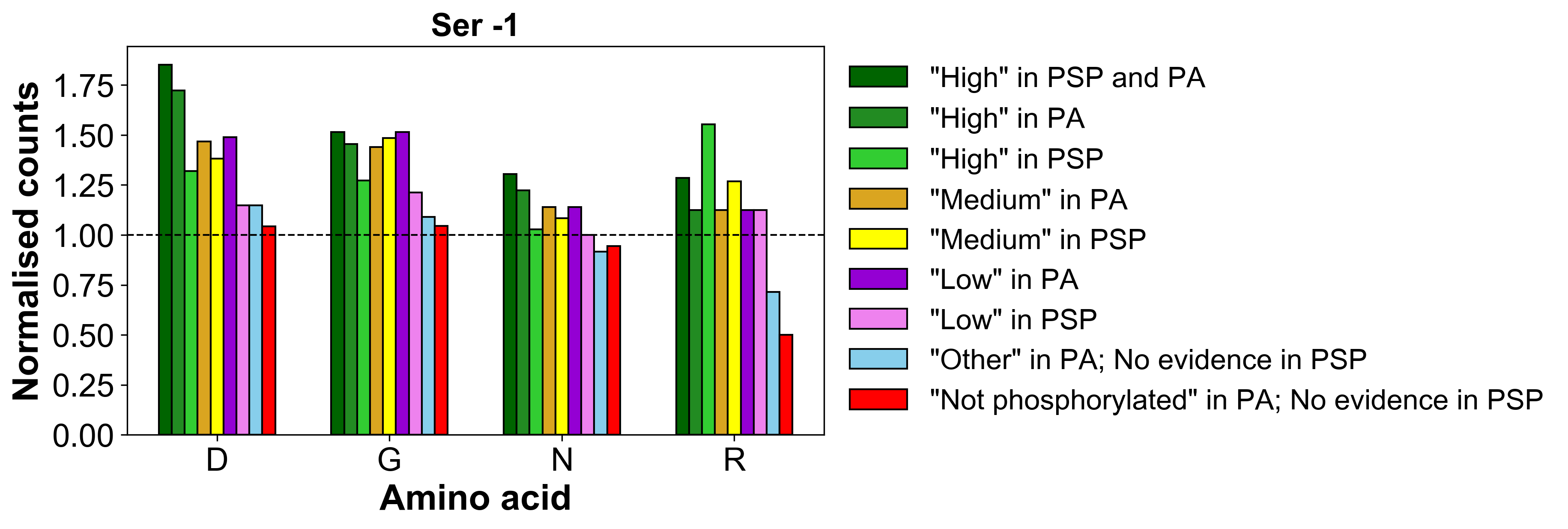

**D**

**C**

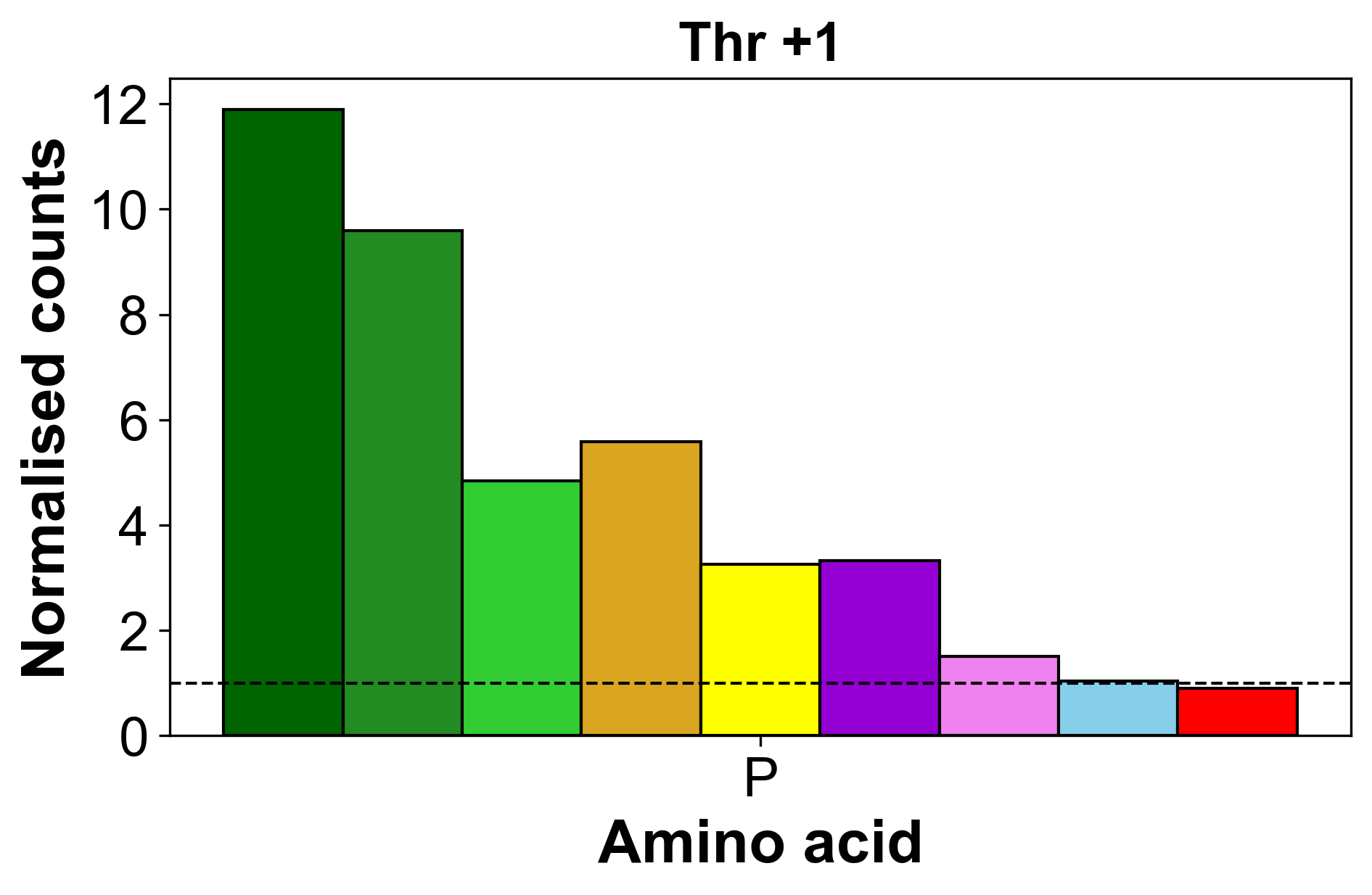

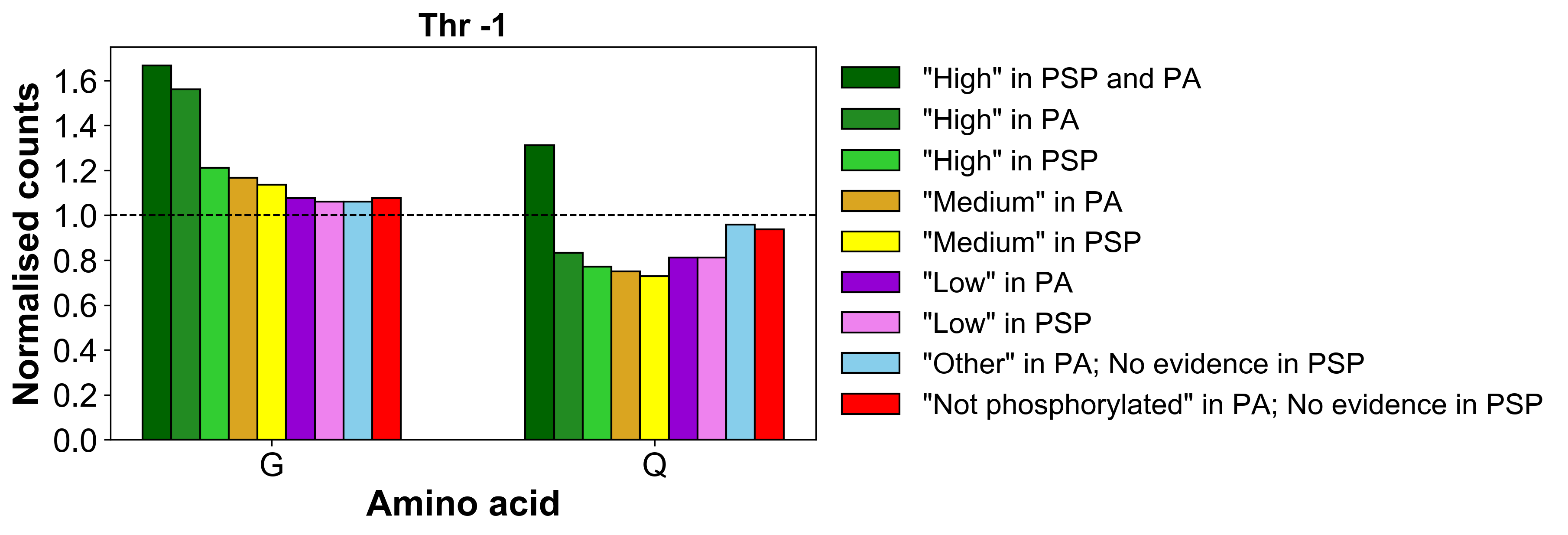

**E**

**F**

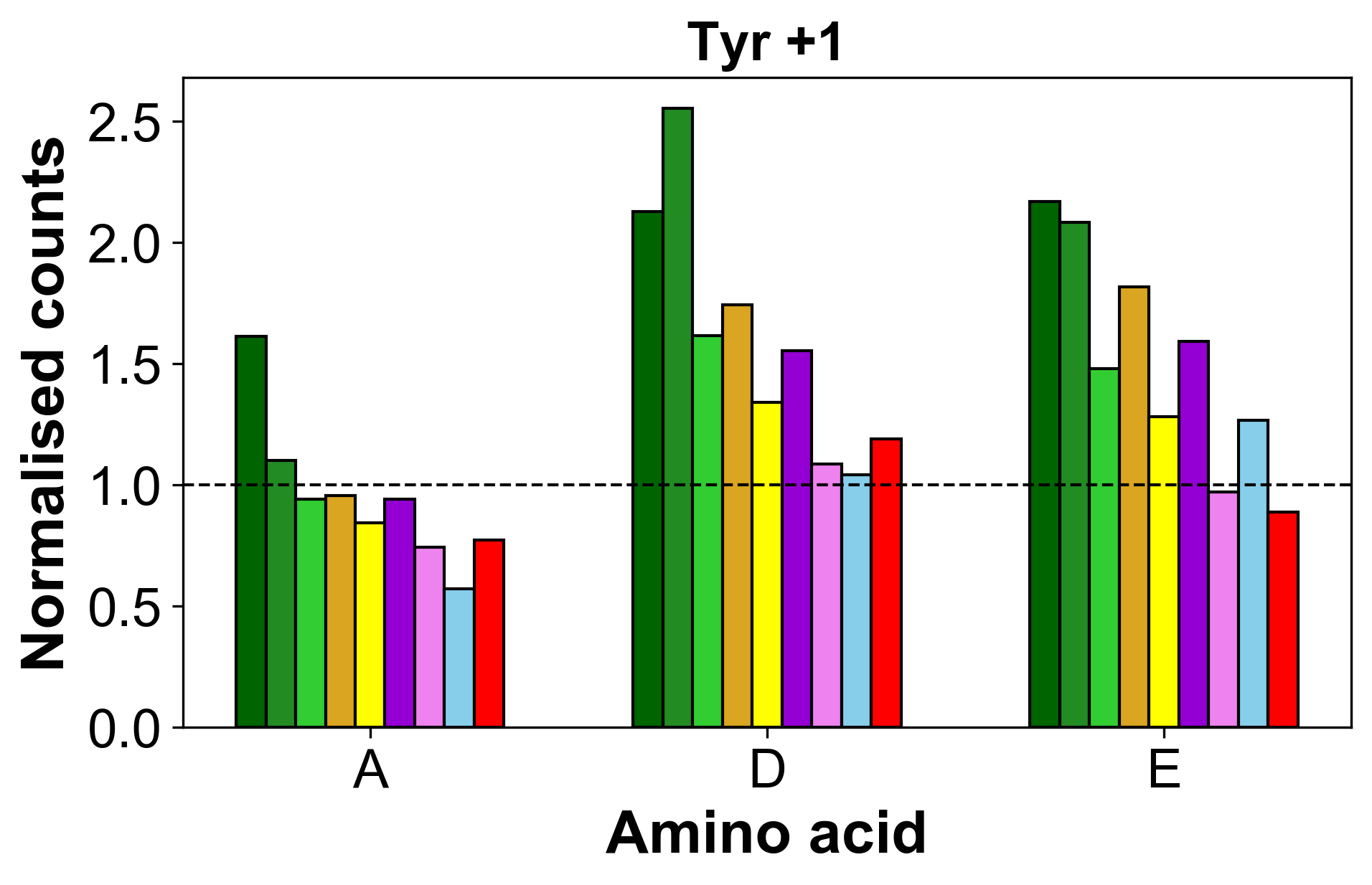

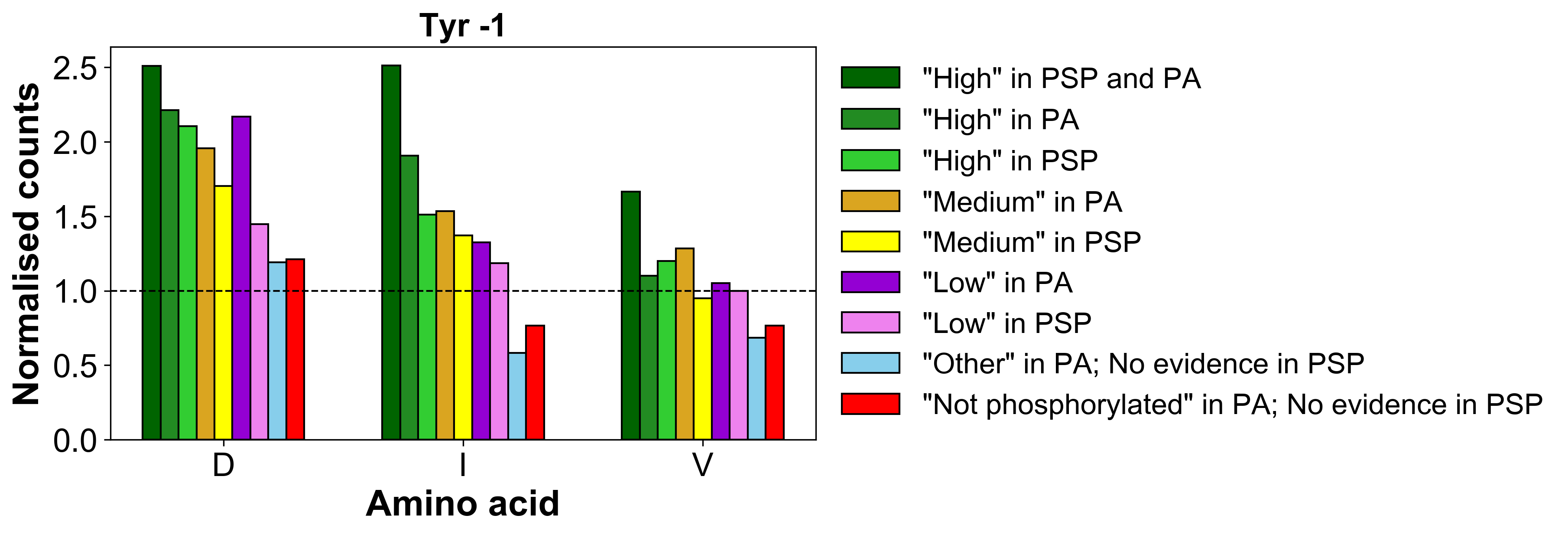

**SI Figure 2.** Normalised counts of proximal amino acids positioned at **(A)** +1 around Ser; **(B)** -1 around Ser; **(C)** +1 around Thr; **(D)** -1 around Thr; **(E)** +1 around Tyr; **(F)** -1 around Tyr sites of various phosphorylation likelihood based on evidence in PSP and PA, which are significantly (Bonferroni corrected p value < 0.01) enriched in the “*High” in PSP and PA*” compared to the “*Not phosphorylated*” set and to the expected amino acid distribution in the human proteome (represented by dotted baseline).
