## Supplementary material for "Profiling the Human Phosphoproteome to Estimate the True Extent of Protein Phosphorylation": SI_Fig3_significant_group_count

**B**

**A**

**C**

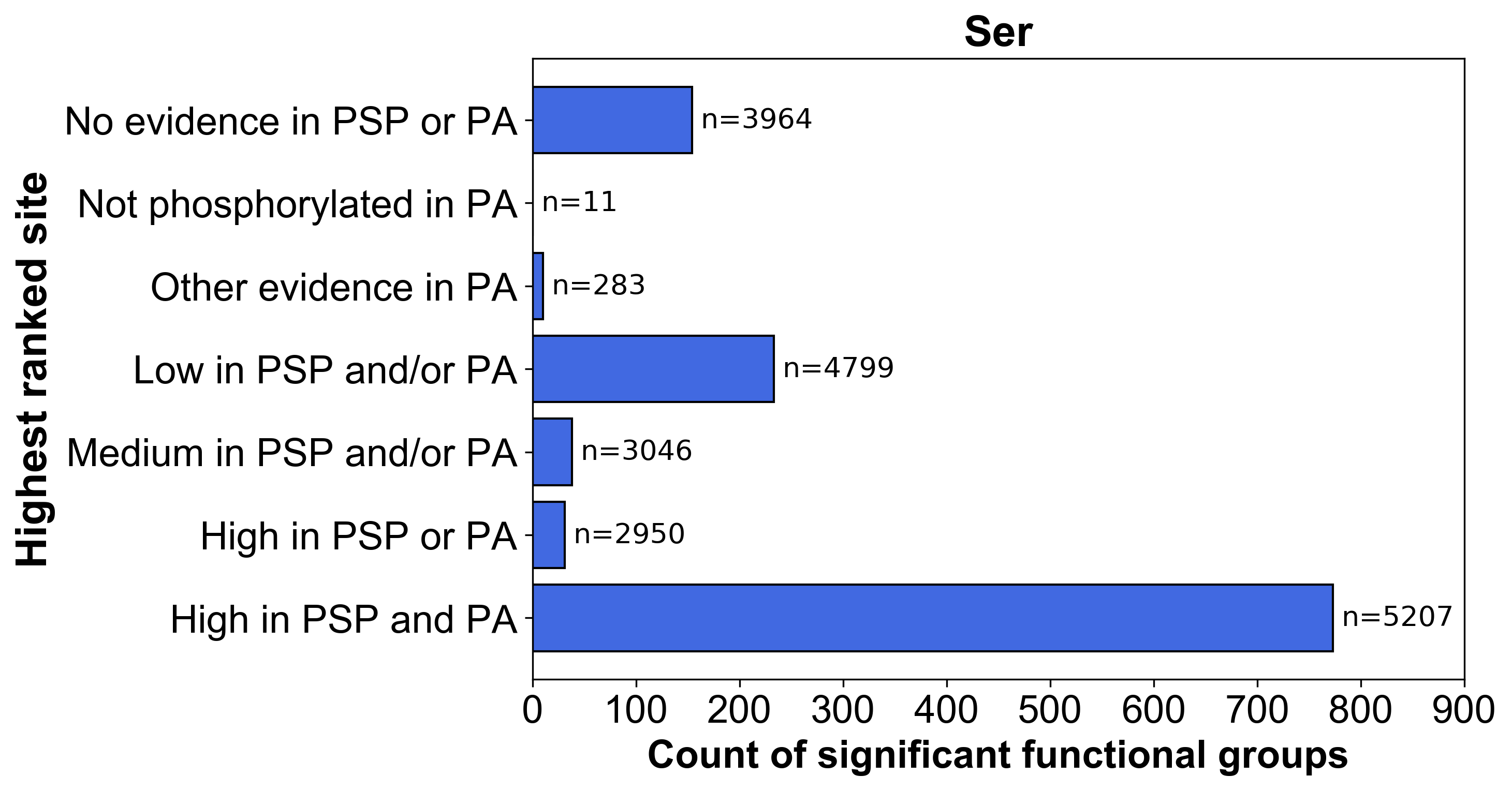

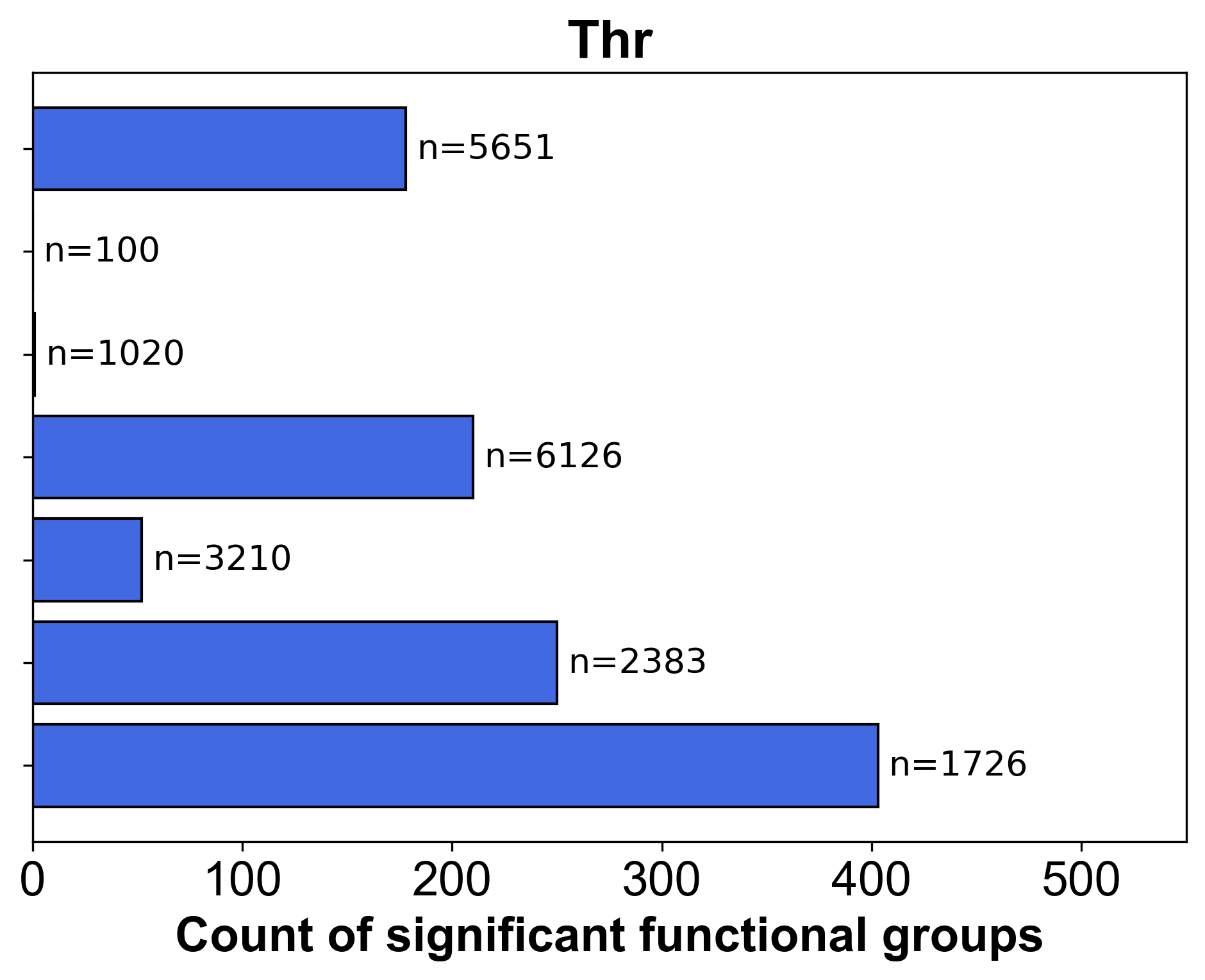

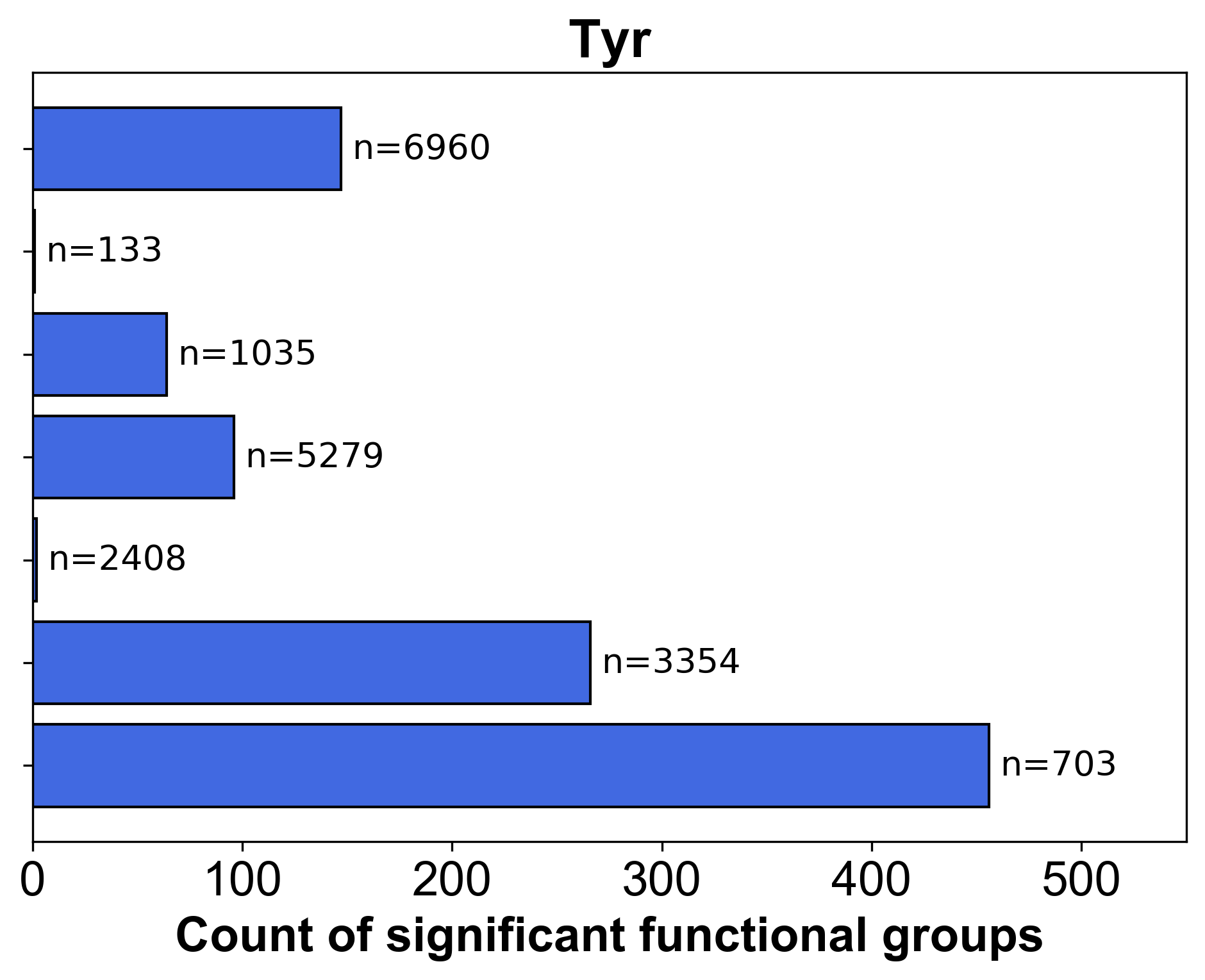

**SI Figure 3.** Count of significant (Benjamini–Hochberg adj. p value <0.05) functional groups identified in DAVID for protein sets containing different highest ranked **(A)** Ser, **(B)** Thr, **(C)** Tyr sites based on phosphorylation likelihood sets in PSP and PA. The number of proteins in each set is presented by n.
