## Supplementary material for "Profiling the Human Phosphoproteome to Estimate the True Extent of Protein Phosphorylation": SI_Fig4_all_sets_in_functional_analysis

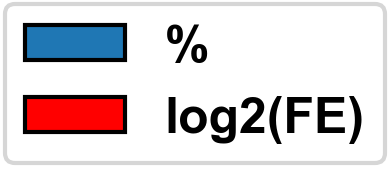

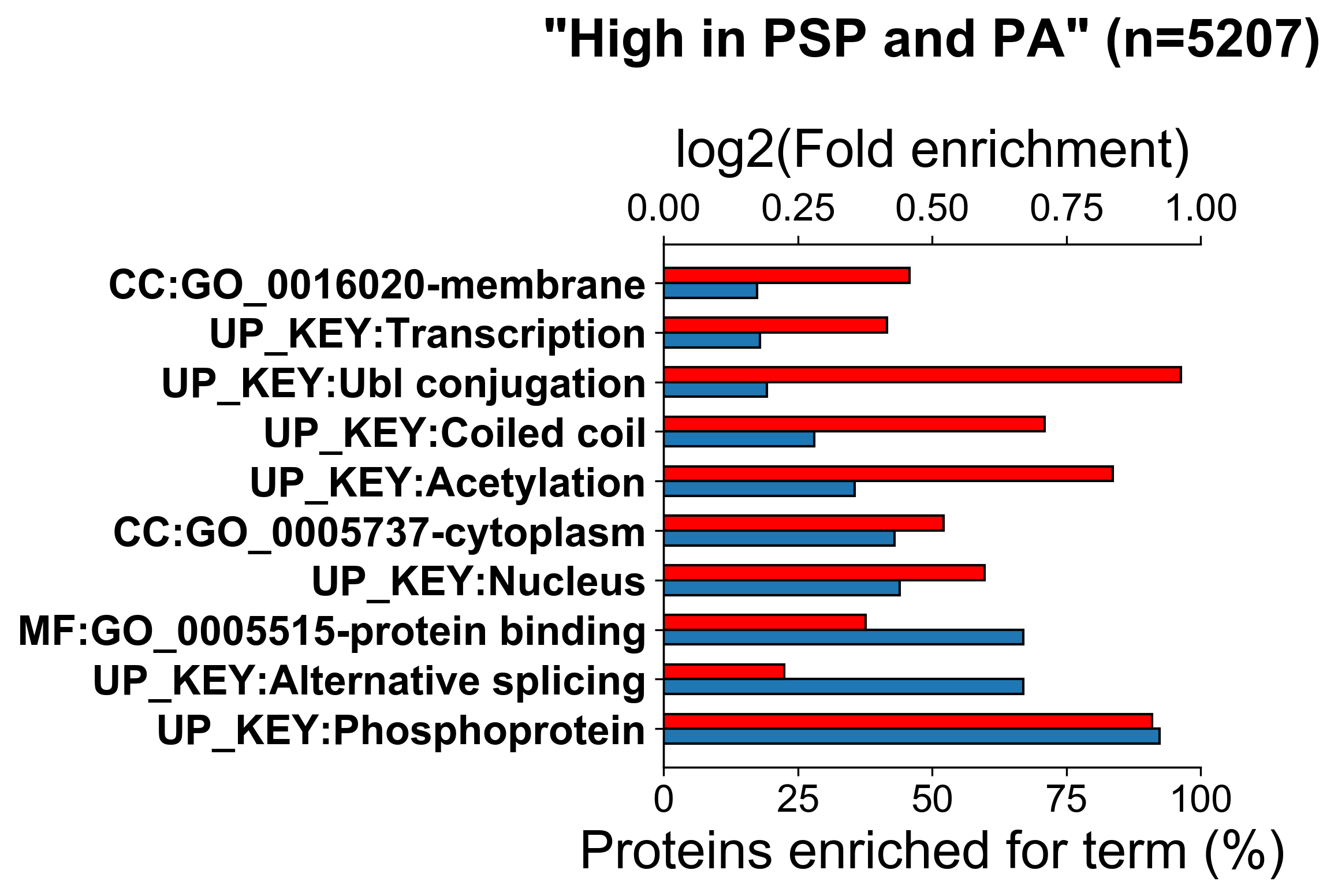

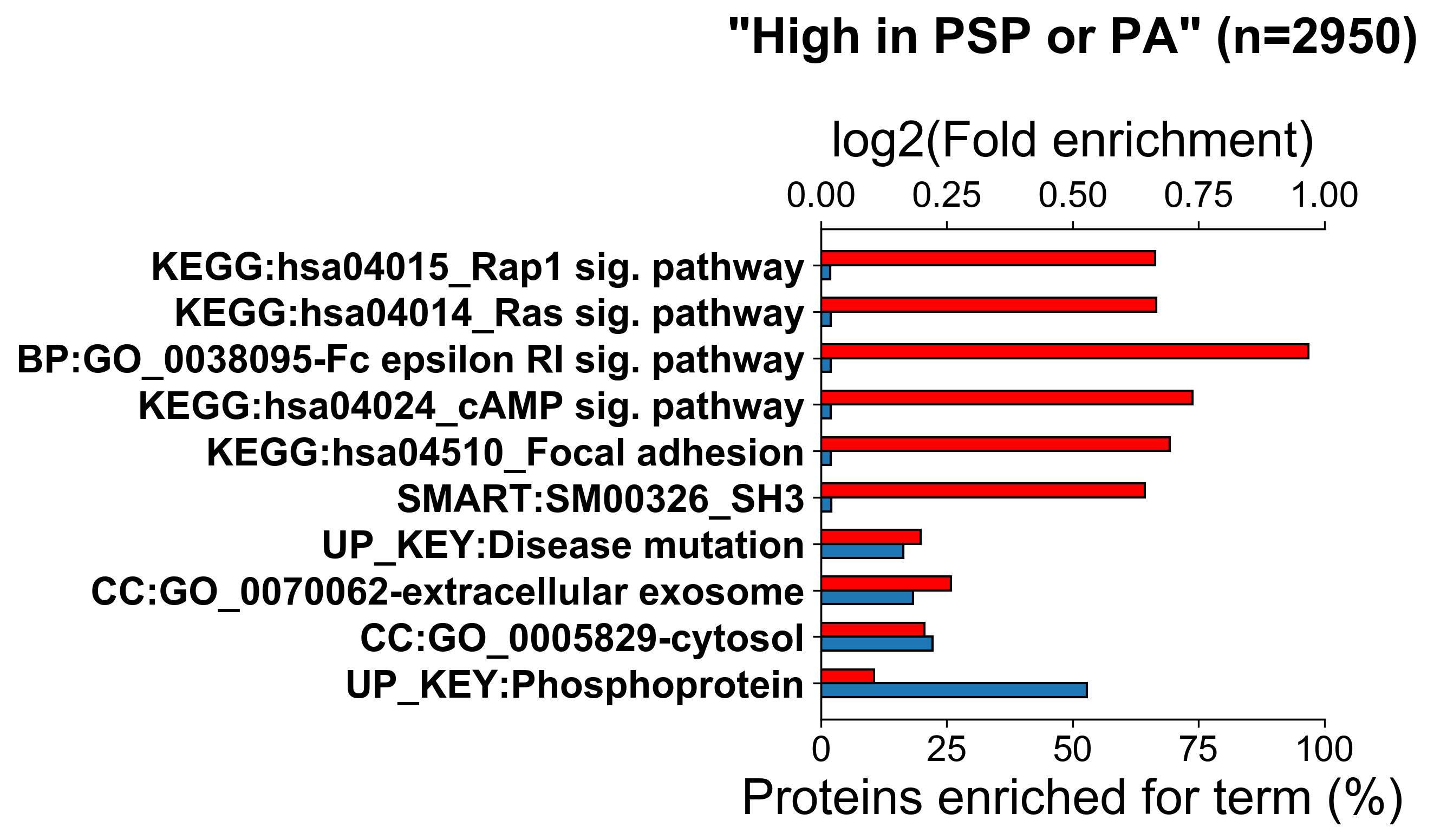

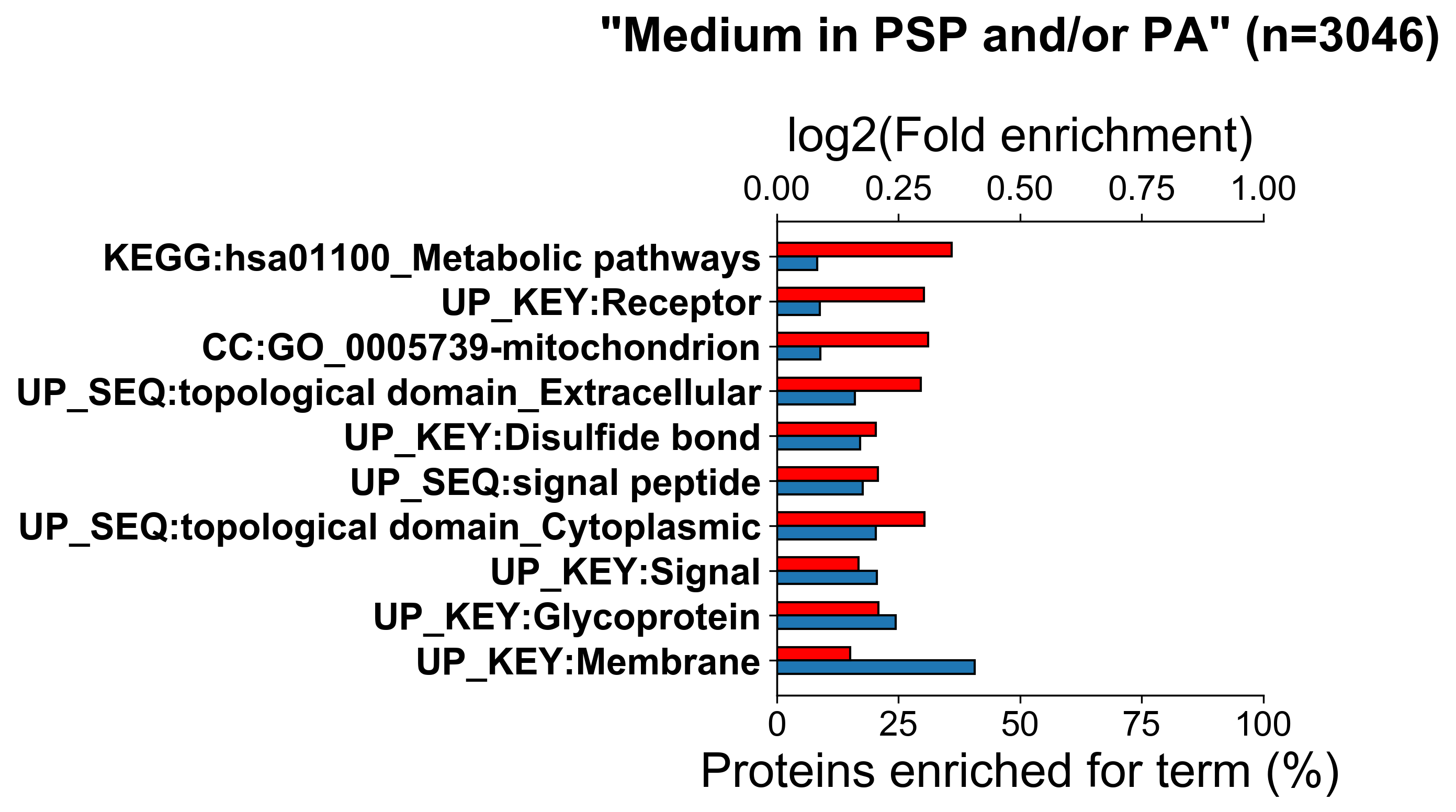

**A**

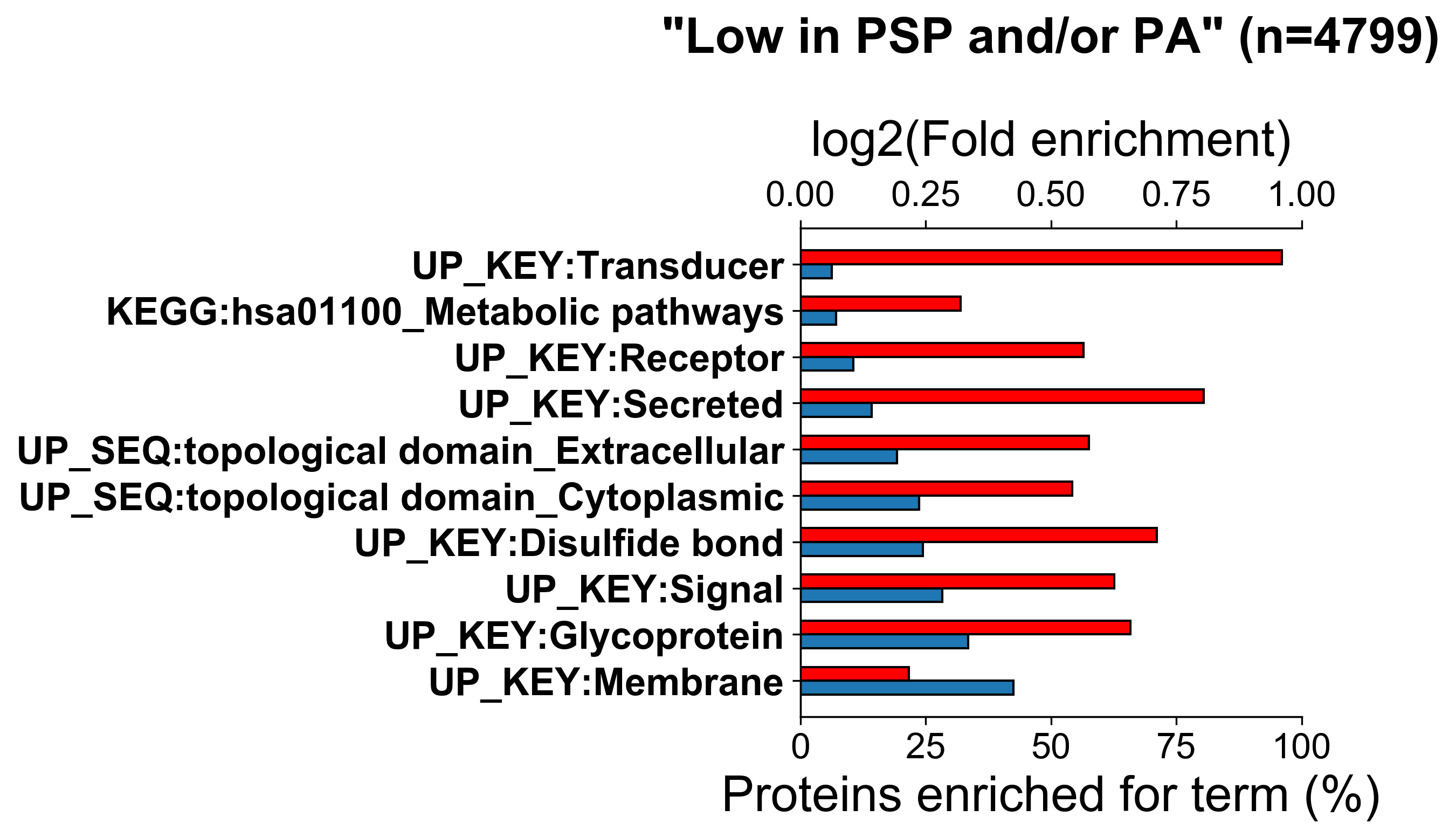

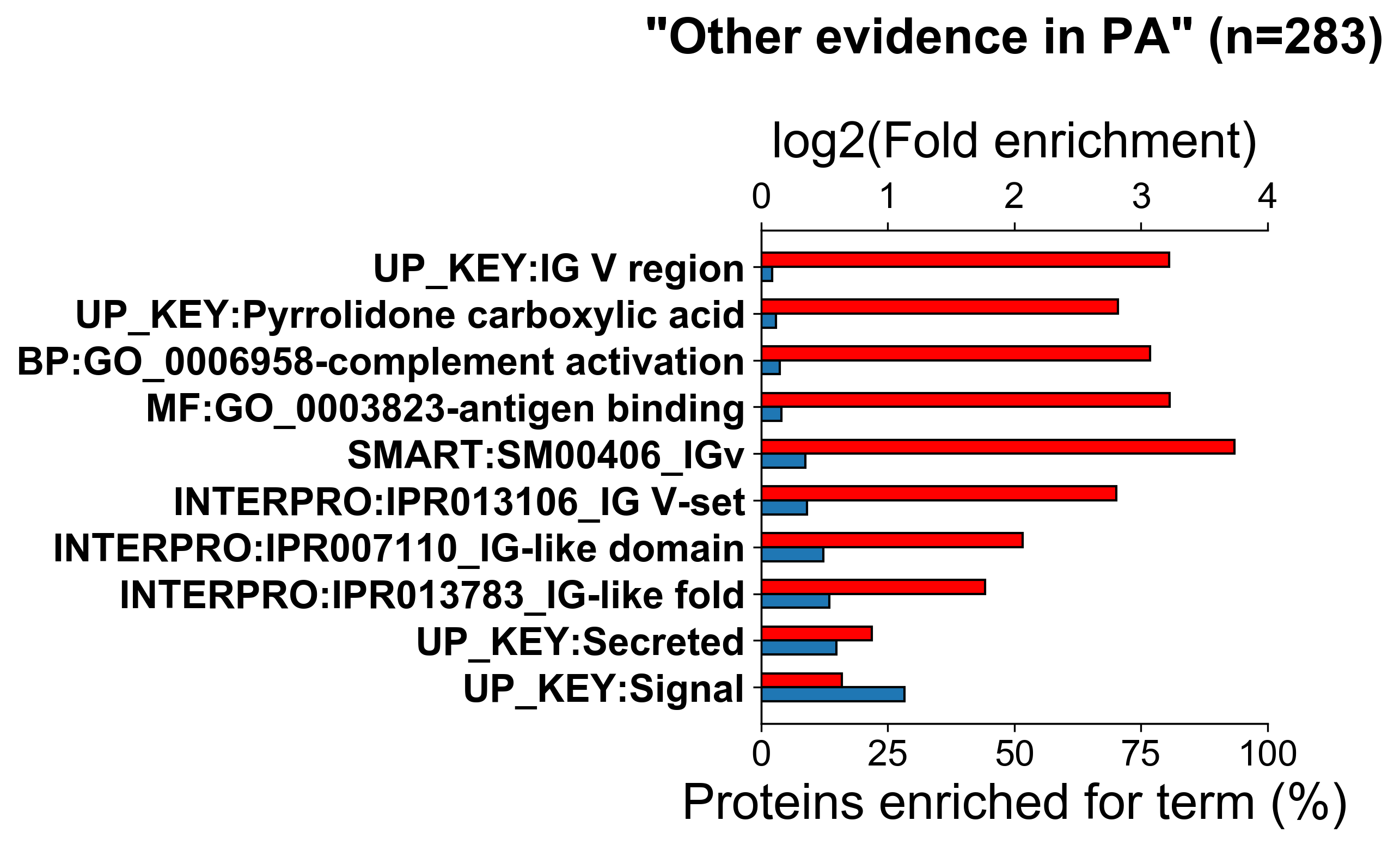

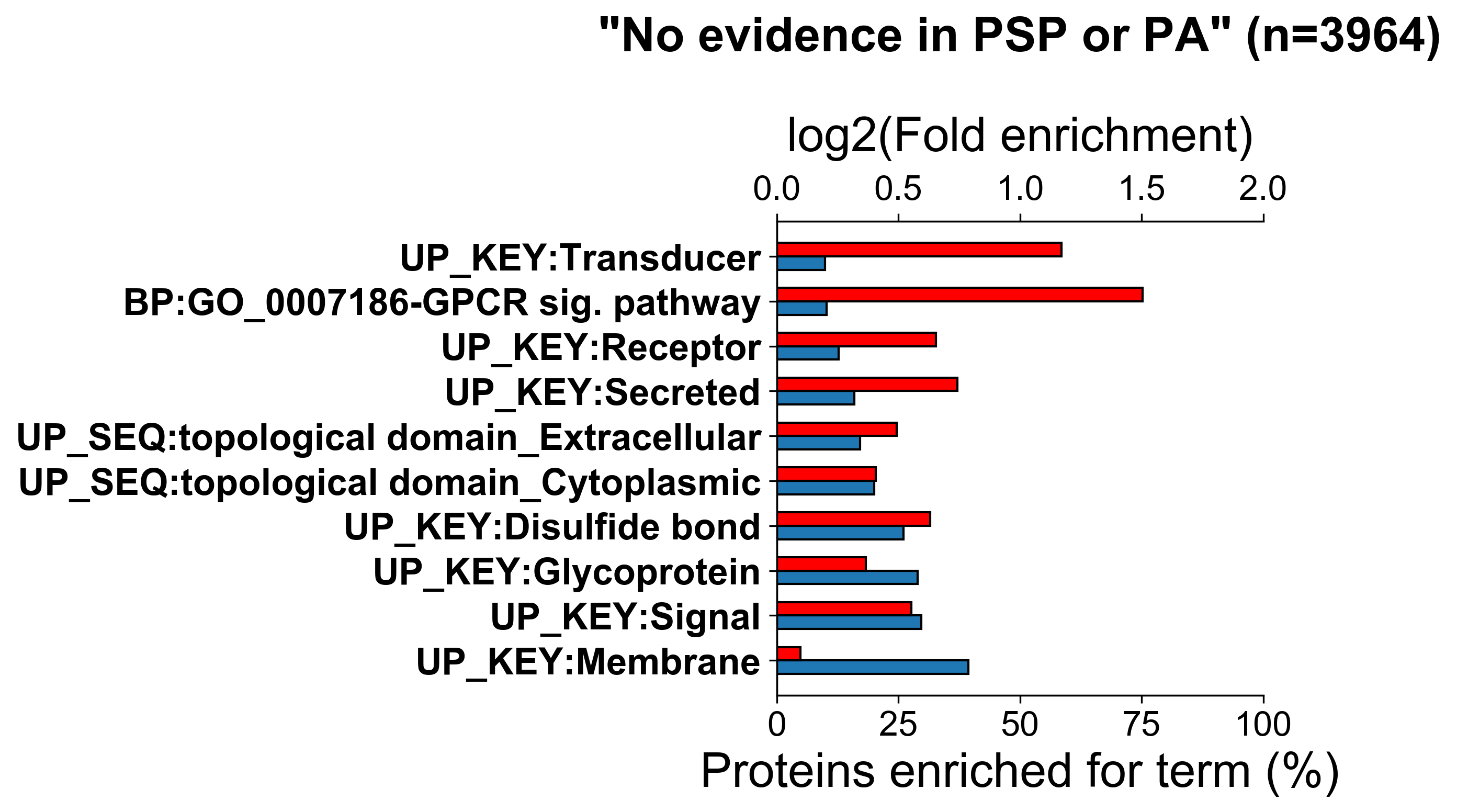

**B**

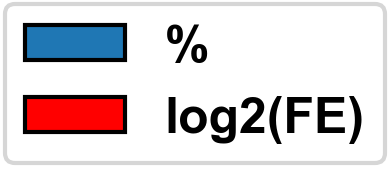

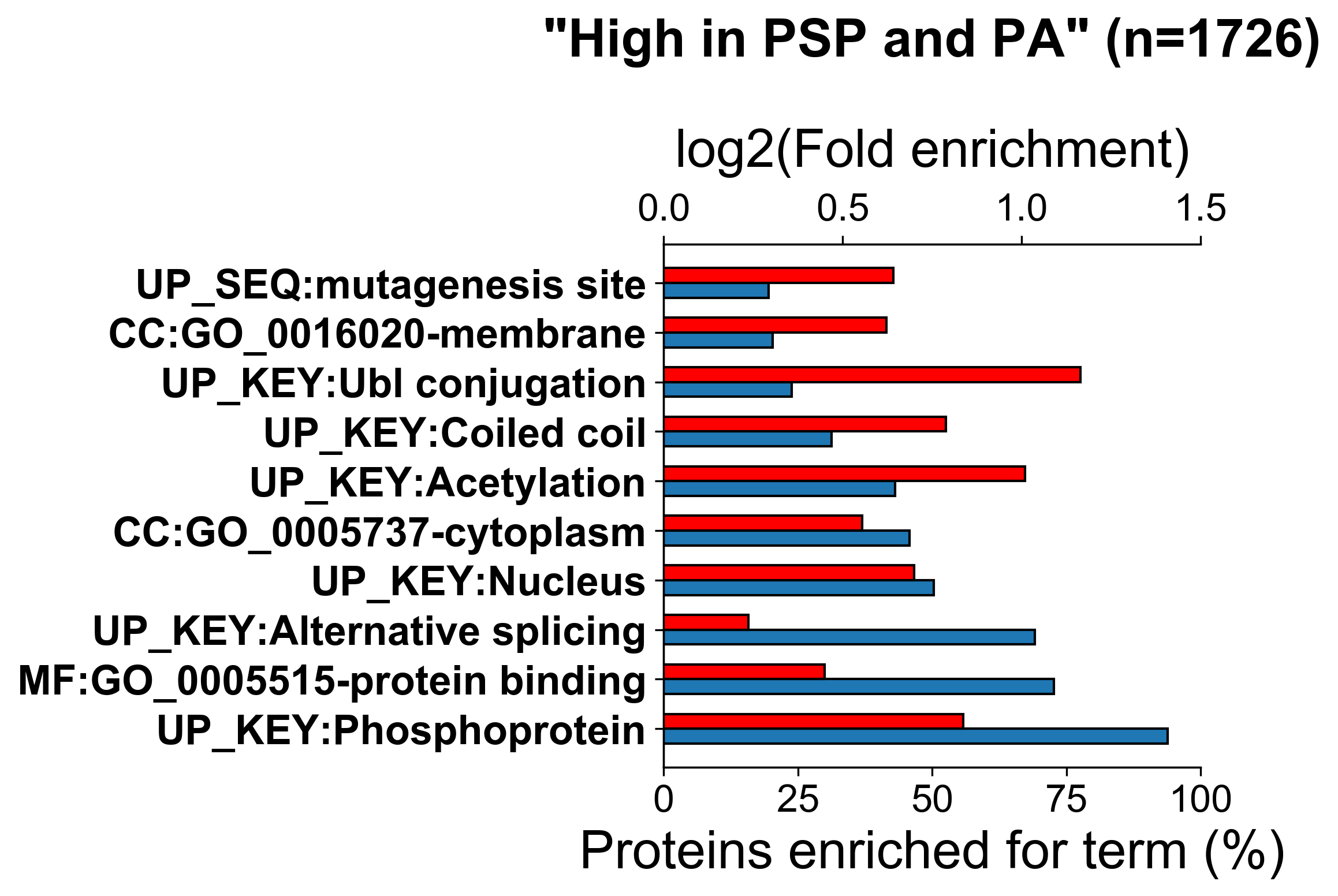

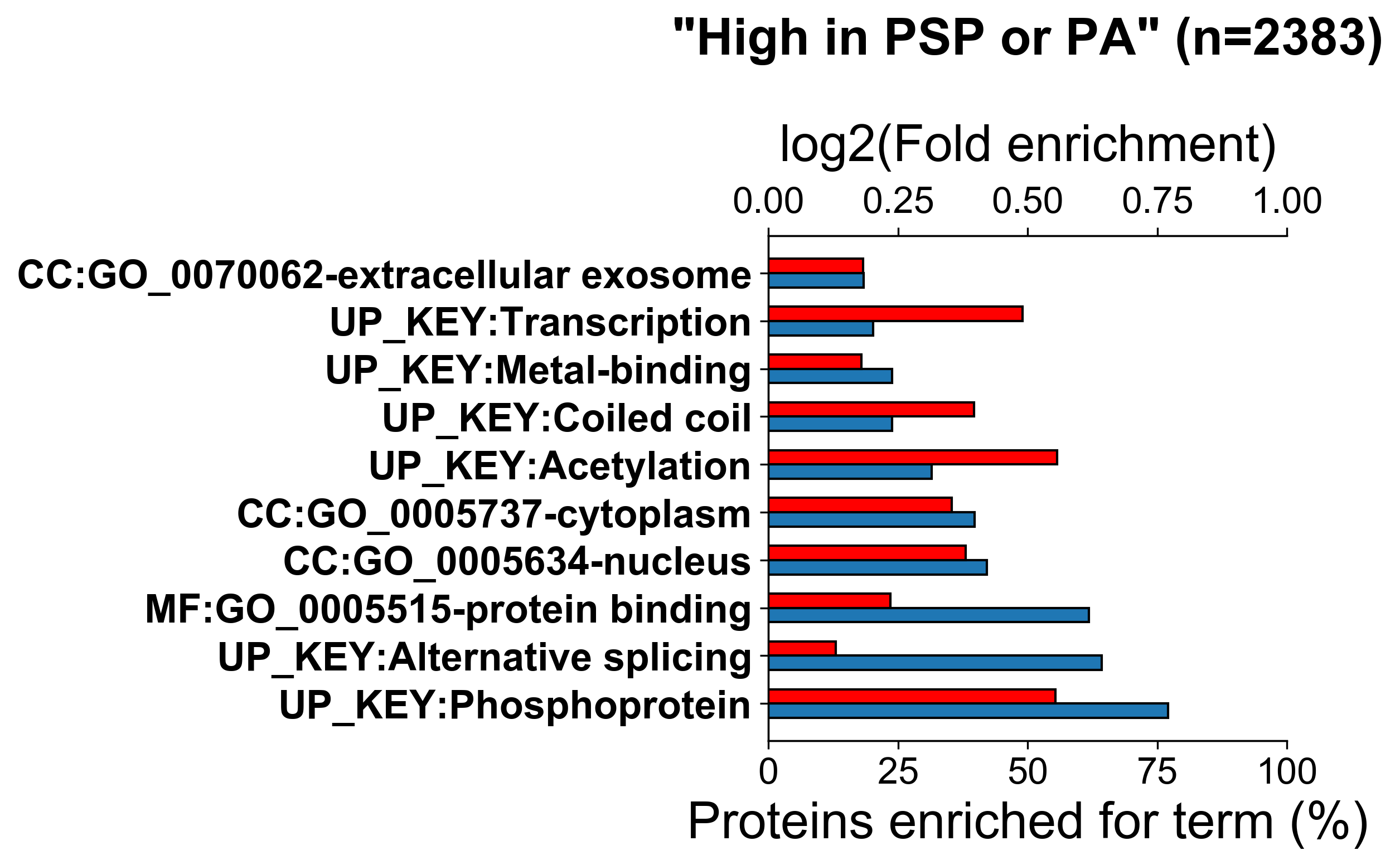

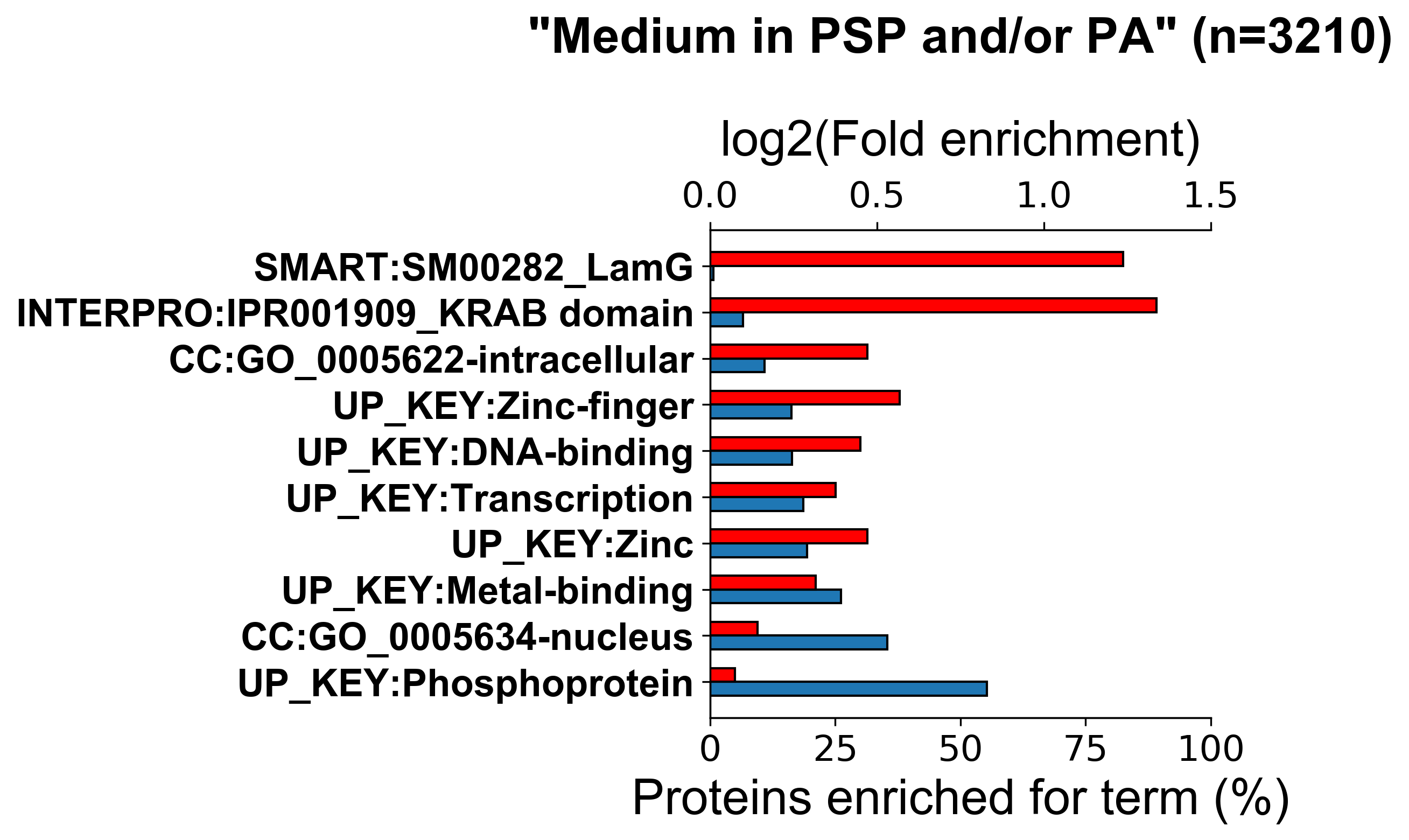

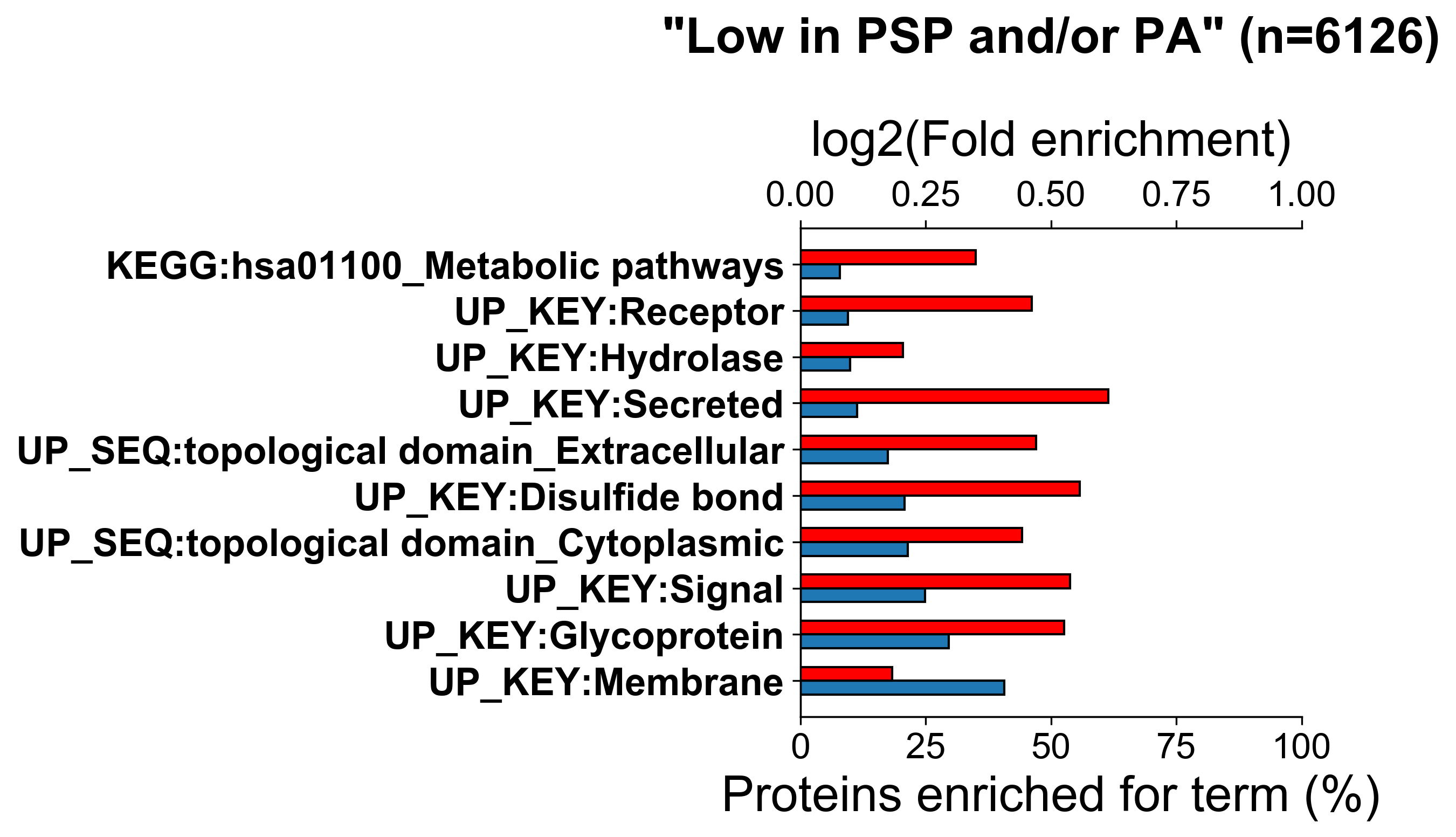

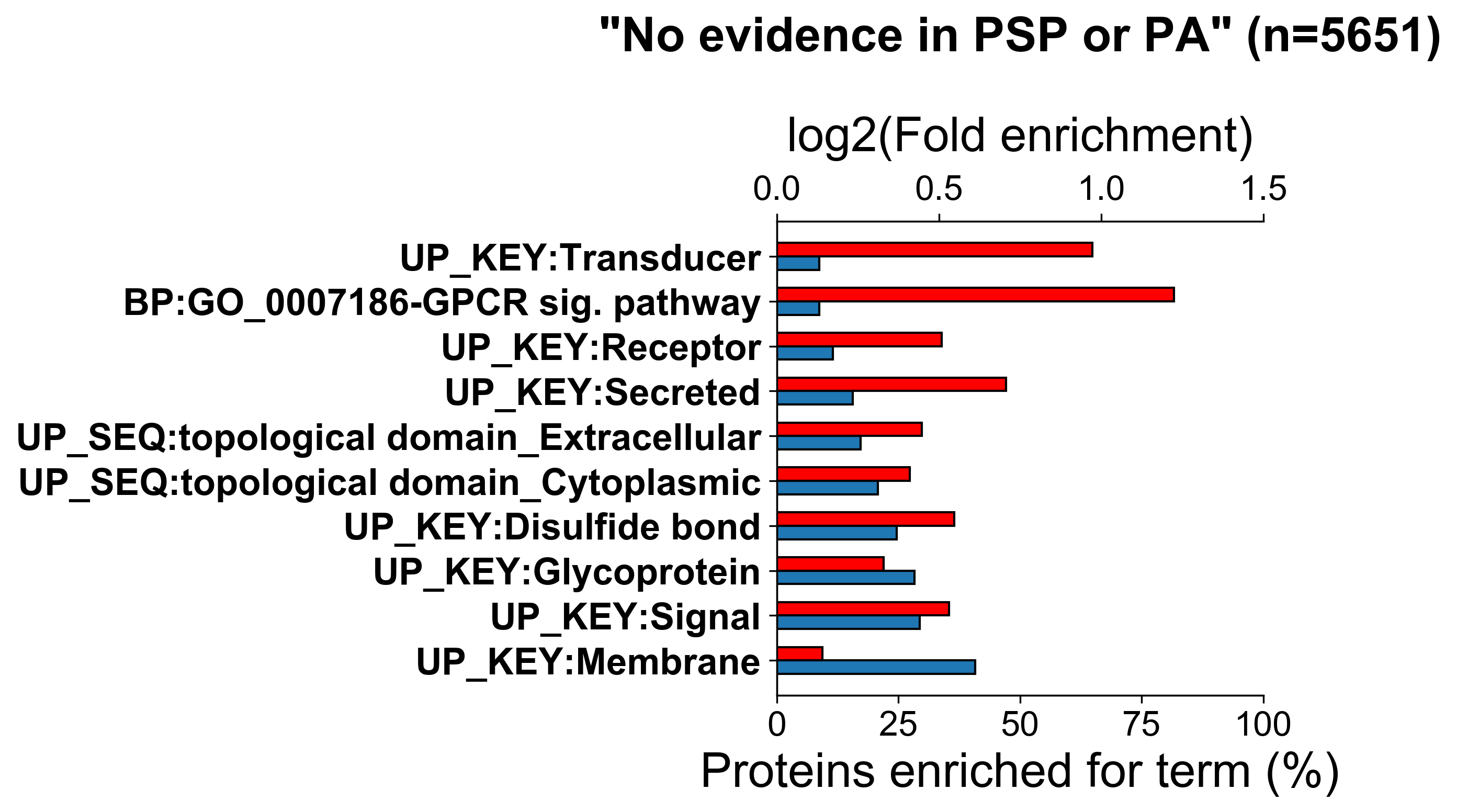

**C**

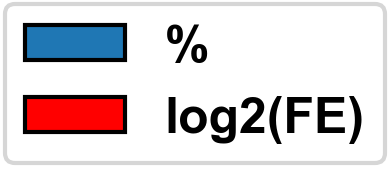

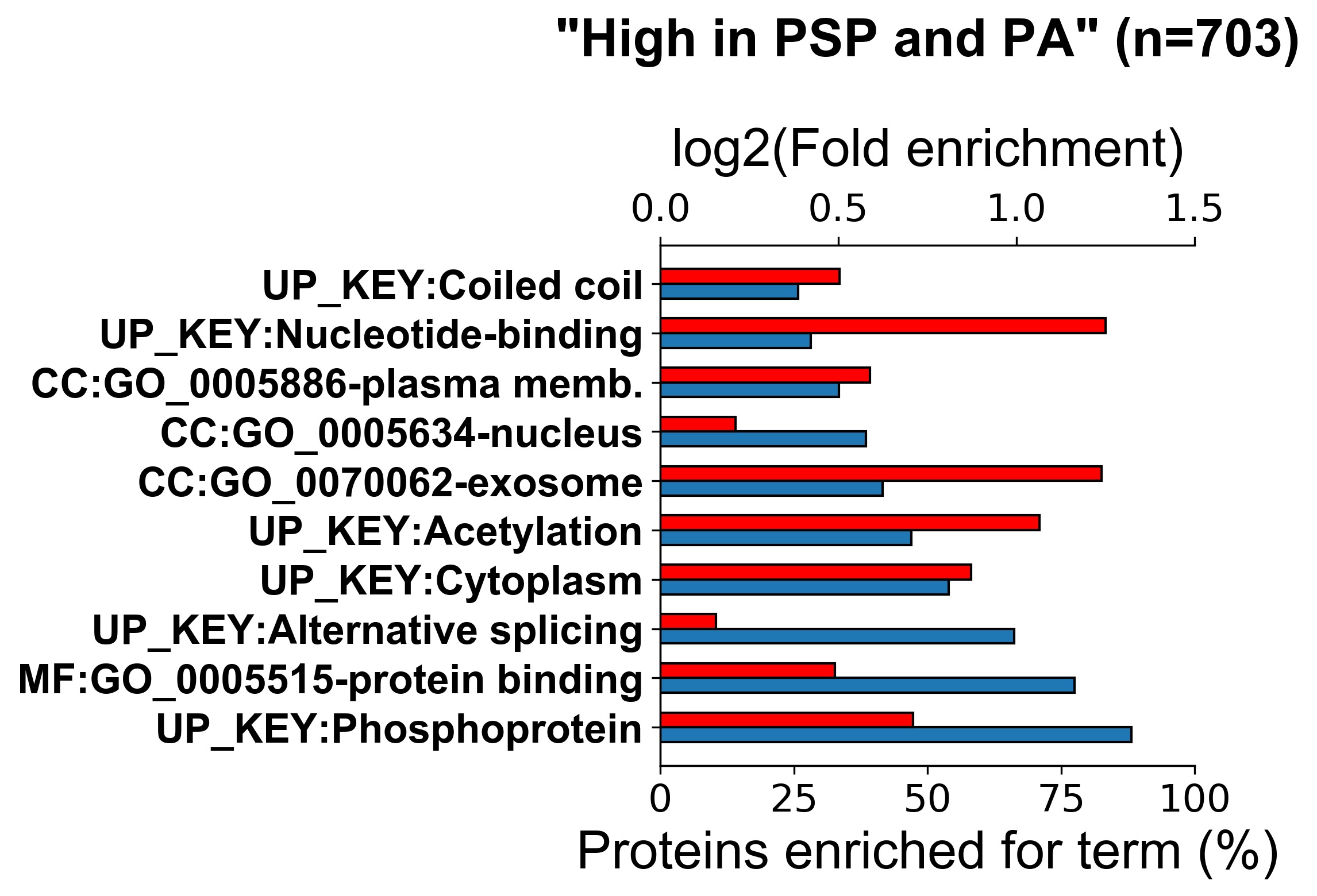

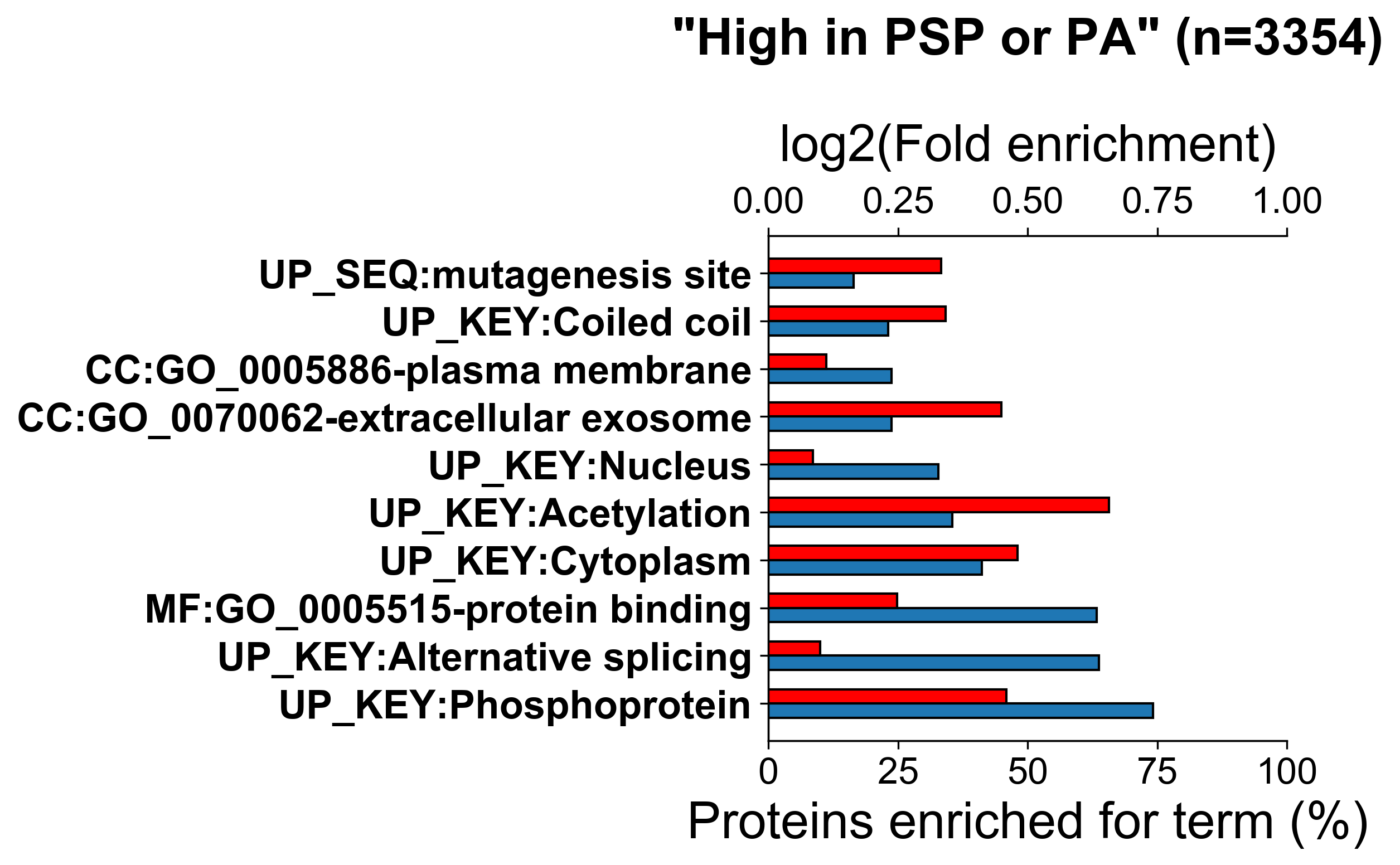

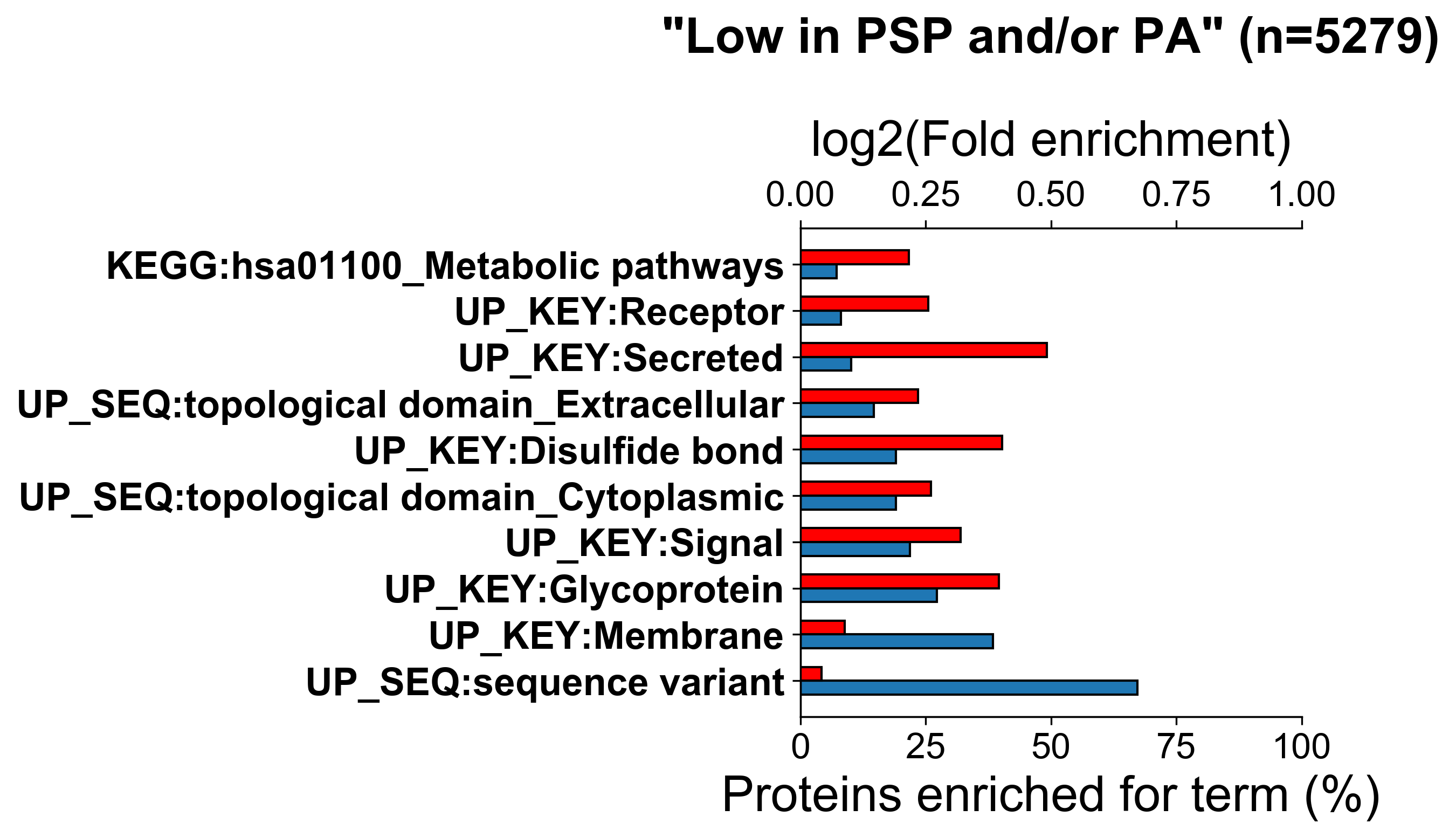

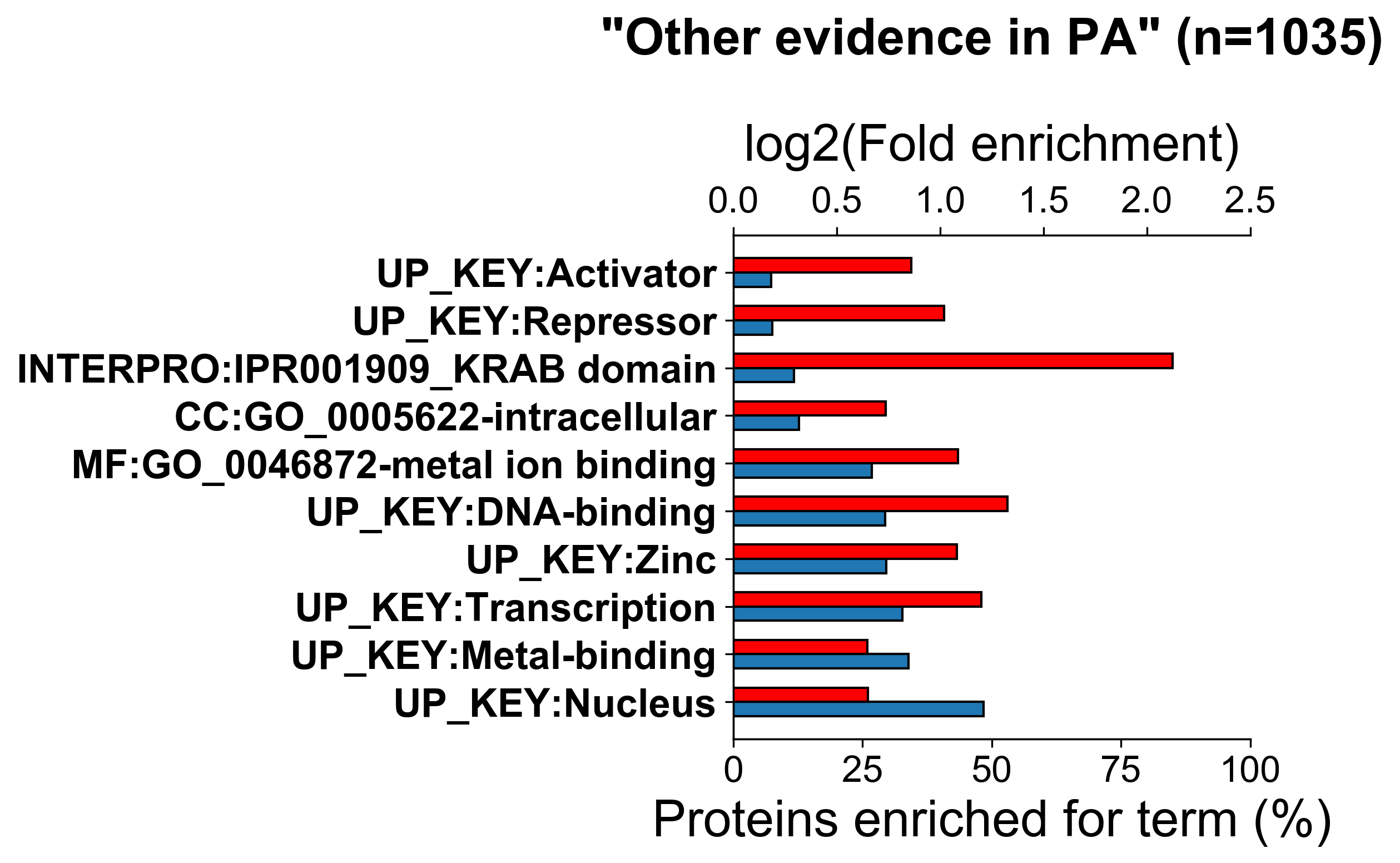

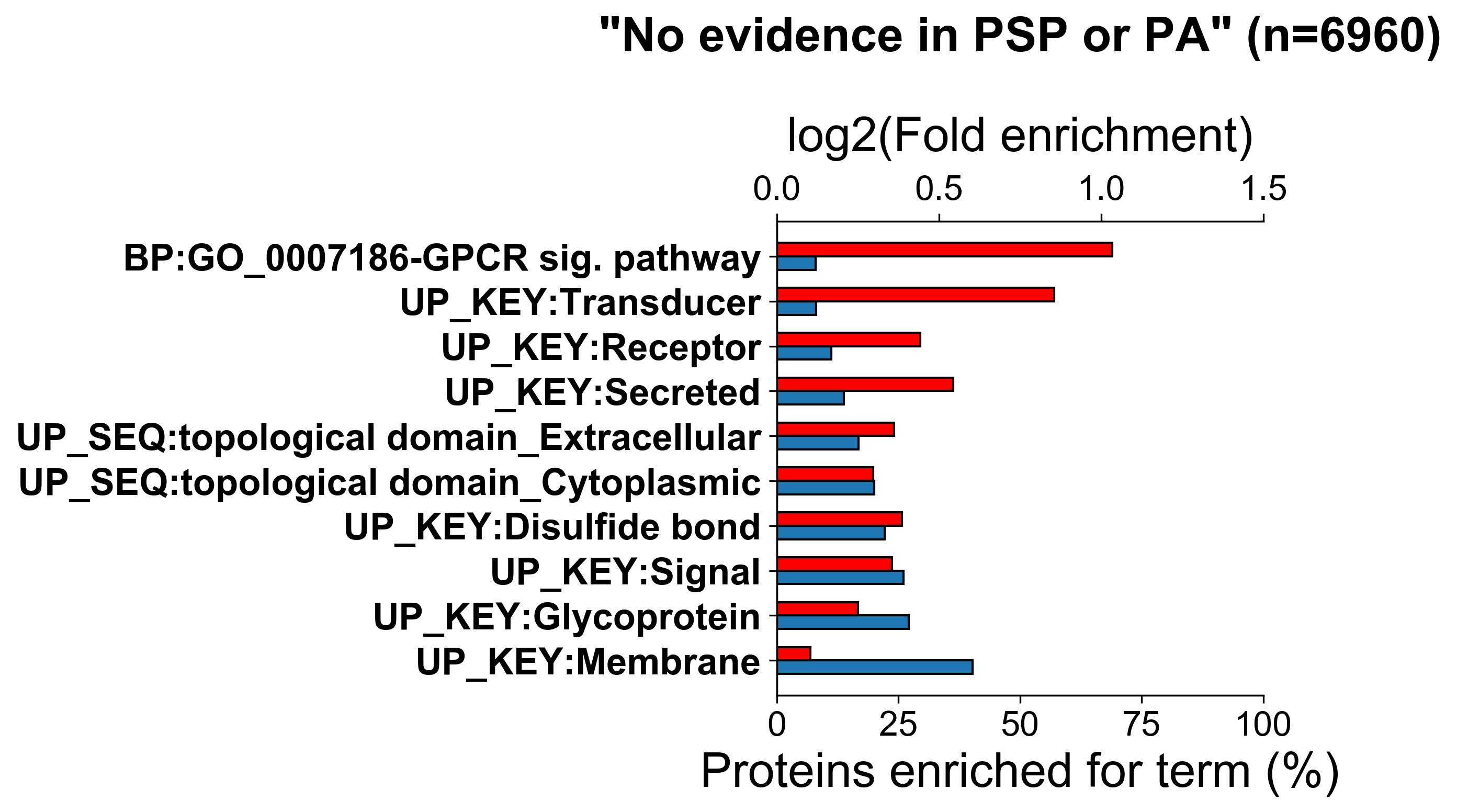

**SI Figure 4.** Top 10 functional categories for which protein sets containing various highest ranked **(A)** Ser, **(B)** Thr, **(C)** Tyr sites based on the amount of available phosphorylation evidence were significantly enriched in DAVID (Benjamini–Hochberg corrected p value <0.05). For each protein set, the percentage of proteins (%) enriched for a particular functional category is given as well as the log2(fold enrichment) for that category. The number of proteins in each set is presented by n.
