## Supplementary material for "Profiling the Human Phosphoproteome to Estimate the True Extent of Protein Phosphorylation": SI_Tab3_proteomes_table

| Organism name from Uniprot | Proteome ID | Organism ID | BUSCO score | Gene count |
| --- | --- | --- | --- | --- |
| *Pan troglodytes* (Chimpanzee) | UP000002277 | 9598 | C:98.3%[S:48.4%,D:49.9%],F:0.5%,M:1.1%,n:6192 | 23003 |
| *Pan paniscus* (Pygmy chimpanzee) | UP000240080 | 9597 | C:97.4%[S:54.4%,D:43%],F:1.4%,M:1.1%,n:6192 | 21211 |
| *Gorilla gorilla gorilla* (Western lowland gorilla) | UP000001519 | 9595 | C:97.3%[S:52%,D:45.3%],F:1.8%,M:1%,n:6192 | 21787 |
| *Pongo abelii* (Sumatran orangutan) | UP000001595 | 9601 | C:94.5%[S:88.5%,D:6%],F:4.2%,M:1.3%,n:6192 | 21992 |
| *Nomascus leucogenys* (Northern white-cheeked gibbon) | UP000001073 | 61853 | C:96.3%[S:57.7%,D:38.6%],F:2.6%,M:1.1%,n:6192 | 20753 |
| *Macaca mulatta* (Rhesus macaque) | UP000006718 | 9544 | C:89.6%[S:47.8%,D:41.8%],F:1.5%,M:8.9%,n:6192 | 21868 |
| *Cercocebus atys* (Sooty mangabey) | UP000233060 | 9531 | C:98%[S:48.3%,D:49.7%],F:1.1%,M:0.9%,n:6192 | 20874 |
| *Rhinopithecus bieti* (Black snub-nosed monkey) | UP000233180 | 61621 | C:95.1%[S:49.6%,D:45.5%],F:2.9%,M:2%,n:6192 | 20845 |
| *Chlorocebus sabaeus* (Green monkey) | UP000029965 | 60711 | C:95.4%[S:94.4%,D:1%],F:3.4%,M:1.2%,n:6192 | 19136 |
| *Macaca fascicularis* (Crab-eating macaque) | UP000233100 | 9541 | C:98.2%[S:48.8%,D:49.3%],F:1.1%,M:0.8%,n:6192 | 22278 |
| *Papio anubis* (Olive baboon) | UP000028761 | 9555 | C:98.4%[S:51.7%,D:46.7%],F:0.8%,M:0.8%,n:6192 | 21559 |
| *Mandrillus leucophaeus* (Drill) | UP000233140 | 9568 | C:95.5%[S:55.1%,D:40.3%],F:3%,M:1.6%,n:6192 | 20767 |
| *Saimiri boliviensis boliviensis* (Bolivian squirrel monkey) | UP000233220 | 39432 | C:96.3%[S:50.1%,D:46.2%],F:2.2%,M:1.5%,n:6192 | 19356 |
| *Callithrix jacchus* (White-tufted-ear marmoset) | UP000008225 | 9483 | C:97.9%[S:52.7%,D:45.2%],F:1%,M:1.1%,n:6192 | 22587 |
| *Aotus nancymaae* (Ma's night monkey) | UP000233020 | 37293 | C:97.4%[S:50.9%,D:46.5%],F:1.3%,M:1.3%,n:6192 | 20363 |
| *Tarsius syrichta* (Philippine tarsier) | UP000189704 | 1868482 | C:75.6%[S:60%,D:15.6%],F:5.7%,M:18.7%,n:6192 | 19956 |
| *Otolemur garnettii* (Small-eared galago) | UP000005225 | 30611 | C:96.8%[S:94%,D:2.8%],F:2.1%,M:1.1%,n:6192 | 19443 |
| *Propithecus coquereli* (Coquerel's sifaka) | UP000233160 | 379532 | C:92%[S:63.6%,D:28.4%],F:4.1%,M:3.9%,n:6192 | 17876 |
| *Ictidomys tridecemlineatus* (Thirteen-lined ground squirrel) | UP000005215 | 43179 | C:94.2%[S:71.1%,D:23%],F:2.6%,M:3.2%,n:6192 | 18440 |
| *Cavia porcellus* (Guinea pig) | UP000005447 | 10141 | C:94.2%[S:70.3%,D:23.9%],F:2.9%,M:2.9%,n:6192 | 18247 |
| *Mus musculus* (Mouse) | UP000000589 | 10090 | C:99.7%[S:52.1%,D:47.6%],F:0.2%,M:0.1%,n:6192 | 21982 |
| *Oryctolagus cuniculus* (Rabbit) | UP000001811 | 9986 | C:91.2%[S:84.3%,D:6.9%],F:4.6%,M:4.1%,n:6192 | 21178 |
| *Cricetulus griseus* (Chinese hamster) | UP000001075 | 10029 | C:63.7%[S:63%,D:0.7%],F:19.2%,M:17.1%,n:6192 | 23874 |
| *Fukomys damarensis* (Damaraland mole rat) | UP000028990 | 885580 | C:82.4%[S:81.9%,D:0.5%],F:7.7%,M:9.9%,n:6192 | 20401 |
| *Mesocricetus auratus* (Golden hamster) | UP000189706 | 10036 | C:77.6%[S:54.7%,D:22.9%],F:1.9%,M:20.5%,n:6192 | 20418 |
| *Dipodomys ordii* (Ord's kangaroo rat) | UP000081671 | 10020 | C:93.7%[S:66.1%,D:27.6%],F:5.4%,M:0.9%,n:6192 | 19730 |
| *Heterocephalus glaber* (Naked mole rat) | UP000006813 | 10181 | C:87%[S:85.5%,D:1.5%],F:6.7%,M:6.2%,n:6192 | 21445 |
| *Vombatus ursinus* (Common wombat) | UP000314987 | 29139 | C:96.6%[S:56.2%,D:40.4%],F:1.8%,M:1.6%,n:4104 | 19872 |
| *Myotis lucifugus* (Little brown bat) | UP000001074 | 59463 | C:95.1%[S:90%,D:5.1%],F:3.5%,M:1.4%,n:6253 | 19655 |
| *Canis lupus familiaris* (Dog) | UP000002254 | 9615 | C:97%[S:45.5%,D:51.5%],F:1.7%,M:1.3%,n:6253 | 20624 |
| *Capra hircus* (Goat) | UP000291000 | 9925 | C:98%[S:57.9%,D:40.1%],F:1.2%,M:0.8%,n:6253 | 21149 |
| *Ovis aries* (Sheep) | UP000002356 | 9940 | C:98%[S:86.8%,D:11.2%],F:1.4%,M:0.5%,n:6253 | 21212 |
| *Sus scrofa* (Pig) | UP000008227 | 9823 | C:95.3%[S:49%,D:46.3%],F:2.4%,M:2.3%,n:6253 | 22130 |
| *Felis catus* (Cat) | UP000011712 | 9685 | C:97%[S:67%,D:30%],F:1.4%,M:1.6%,n:6253 | 19645 |
| *Ailuropoda melanoleuca* (Giant panda) | UP000008912 | 9646 | C:97.4%[S:91.7%,D:5.8%],F:2%,M:0.6%,n:6253 | 19332 |
| *Pteropus alecto* (Black flying fox) | UP000010552 | 9402 | C:84.2%[S:83.7%,D:0.5%],F:9%,M:6.8%,n:6253 | 19520 |
| *Erinaceus europaeus* (Western European hedgehog) | UP000079721 | 9365 | C:95.2%[S:66.7%,D:28.4%],F:3.6%,M:1.2%,n:6253 | 19242 |
| *Equus caballus* (Horse) | UP000002281 | 9796 | C:97.8%[S:57.5%,D:40.3%],F:1.3%,M:0.9%,n:6253 | 20845 |
| *Bos taurus* (Bovine) | UP000009136 | 9913 | C:98.2%[S:34.4%,D:63.8%],F:1.2%,M:0.6%,n:6253 | 23844 |
| *Mustela putorius furo* (European domestic ferret) | UP000000715 | 9669 | C:96.2%[S:94.9%,D:1.3%],F:2.4%,M:1.4%,n:6253 | 19902 |
| *Lipotes vexillifer* (Yangtze river dolphin) | UP000265300 | 118797 | C:98.5%[S:74.2%,D:24.3%],F:1.1%,M:0.3%,n:6253 | 18846 |
| *Leptonychotes weddellii* (Weddell seal) | UP000245341 | 9713 | C:85%[S:50.5%,D:34.5%],F:14%,M:1%,n:6253 | 13162 |
| *Ursus maritimus* (Polar bear) | UP000261680 | 29073 | C:97.1%[S:71.8%,D:25.3%],F:2.6%,M:0.2%,n:6253 | 19368 |
| *Delphinapterus leucas* (Beluga whale) | UP000248483 | 9749 | C:98.6%[S:44%,D:54.6%],F:1.1%,M:0.4%,n:6253 | 17043 |
| *Odobenus rosmarus divergens* (Pacific walrus) | UP000245340 | 9708 | C:99%[S:64%,D:35%],F:0.9%,M:0.1%,n:6253 | 19331 |
| *Physeter macrocephalus* (Sperm whale) | UP000248484 | 9755 | C:86.3%[S:53.7%,D:32.6%],F:0.8%,M:12.9%,n:6253 | 20100 |
| *Tursiops truncatus* (Atlantic bottle-nosed dolphin) | UP000245320 | 9739 | C:84.1%[S:42.6%,D:41.4%],F:8.4%,M:7.6%,n:6253 | 17075 |
| *Loxodonta africana* (African elephant) | UP000007646 | 9785 | C:97.4%[S:73.3%,D:24.1%],F:1.9%,M:0.6%,n:4104 | 20015 |
| *Trichechus manatus latirostris* (Florida manatee) | UP000248480 | 127582 | C:98.1%[S:55.4%,D:42.7%],F:1.6%,M:0.3%,n:4104 | 19079 |
| *Ornithorhynchus anatinus* (Duckbill platypus) | UP000002279 | 9258 | C:75.7%[S:68%,D:7.7%],F:18.8%,M:5.5%,n:4104 | 21677 |
| *Meleagris gallopavo* (Wild turkey) | UP000001645 | 9103 | C:91.1%[S:81.4%,D:9.7%],F:5.4%,M:3.5%,n:4915 | 14164 |
| *Taeniopygia guttata* (Zebra finch) | UP000007754 | 59729 | C:95.4%[S:91.2%,D:4.3%],F:3.6%,M:0.9%,n:4915 | 17428 |
| *Anas platyrhynchos* (Mallard) | UP000296049 | 8839 | C:79.4%[S:78.5%,D:0.9%],F:7.8%,M:12.8%,n:4915 | 16574 |
| *Dryobates pubescens* (Downy woodpecker) | UP000053875 | 118200 | C:94.6%[S:93.7%,D:1%],F:2.2%,M:3.2%,n:4915 | 13097 |
| *Tinamus guttatus* (White-throated tinamou) | UP000053641 | 94827 | C:89.8%[S:88.5%,D:1.3%],F:5.5%,M:4.7%,n:4915 | 13377 |
| *Amazona aestiva* (Blue-fronted Amazon parrot) | UP000051836 | 12930 | C:86.6%[S:85.4%,D:1.2%],F:7.5%,M:5.9%,n:4915 | 16092 |
| *Calypte anna* (Anna's hummingbird) | UP000054308 | 9244 | C:95.4%[S:94.5%,D:0.9%],F:1.7%,M:2.9%,n:4915 | 13267 |
| *Columba livia* (Rock dove) | UP000053872 | 8932 | C:93%[S:78.5%,D:14.4%],F:5%,M:2.1%,n:4915 | 14619 |
| *Callipepla squamata* (Scaled quail) | UP000198323 | 9009 | C:77.6%[S:75.8%,D:1.8%],F:14.5%,M:7.9%,n:4915 | 16973 |
| *Aptenodytes forsteri* (Emperor penguin) | UP000053286 | 9233 | C:97.9%[S:97%,D:0.9%],F:0.8%,M:1.3%,n:4915 | 13704 |
| *Opisthocomus hoazin* (Hoatzin) | UP000053605 | 30419 | C:95.3%[S:94.7%,D:0.6%],F:2.1%,M:2.6%,n:4915 | 12773 |
| *Egretta garzetta* (Little egret) | UP000053119 | 188379 | C:96.7%[S:96%,D:0.8%],F:1%,M:2.3%,n:4915 | 13489 |
| *Alligator mississippiensis* (American alligator) | UP000050525 | 8496 | C:88.6%[S:67.4%,D:21.2%],F:6.8%,M:4.6%,n:3950 | 24656 |
| *Alligator sinensis* (Chinese alligator) | UP000189705 | 38654 | C:74.4%[S:55.1%,D:19.3%],F:2.4%,M:23.2%,n:3950 | 19111 |
| *Anolis carolinensis* (Green anole) | UP000001646 | 28377 | C:92.1%[S:88.4%,D:3.7%],F:5.3%,M:2.6%,n:3950 | 18525 |
| *Pelodiscus sinensis* (Chinese softshell turtle) | UP000007267 | 13735 | C:93.5%[S:79.1%,D:14.4%],F:4.9%,M:1.6%,n:3950 | 18109 |
| *Salmo salar* (Atlantic salmon) | UP000087266 | 8030 | C:97.9%[S:23.9%,D:74%],F:1.5%,M:0.6%,n:4584 | 47717 |
| *Oncorhynchus mykiss* (Rainbow trout) | UP000193380 | 8022 | C:77.1%[S:46.4%,D:30.7%],F:10.9%,M:12%,n:4584 | 46447 |
| *Gasterosteus aculeatus* (Three-spined stickleback) | UP000007635 | 69293 | C:97.5%[S:74.5%,D:23%],F:1.9%,M:0.6%,n:4584 | 20665 |
| *Seriola dumerili* (Greater amberjack) | UP000261420 | 41447 | C:97.6%[S:69.5%,D:28.1%],F:1.5%,M:0.8%,n:4584 | 23238 |
| *Takifugu rubripes* (Japanese pufferfish) | UP000005226 | 31033 | C:95.1%[S:65.4%,D:29.7%],F:2.7%,M:2.2%,n:4584 | 20591 |
| *Xenopus laevis* (African clawed frog) | UP000186698 | 8355 | C:95.6%[S:41.5%,D:54.1%],F:1.6%,M:2.8%,n:3950 | 43235 |
| *Xenopus tropicalis* (Western clawed frog) | UP000008143 | 8364 | C:62.9%[S:37.4%,D:25.4%],F:2.4%,M:34.8%,n:3950 | 35973 |
| *Daphnia pulex* (Water flea) | UP000000305 | 6669 | C:96.1%[S:93.8%,D:2.3%],F:1.4%,M:2.5%,n:1066 | 30118 |
| *Tribolium castaneum* (Red flour beetle) | UP000007266 | 7070 | C:98.6%[S:91.5%,D:7.1%],F:1.2%,M:0.2%,n:1658 | 16568 |
| *Bombyx mori* (Silk moth) | UP000005204 | 7091 | C:89.9%[S:89.4%,D:0.5%],F:6.8%,M:3.3%,n:1658 | 14773 |
| *Anopheles darlingi* (Mosquito) | UP000000673 | 43151 | C:89.4%[S:89%,D:0.4%],F:2.6%,M:7.9%,n:2799 | 10447 |
| *Drosophila melanogaster* (Fruit fly) | UP000000803 | 7227 | C:99.3%[S:38.2%,D:61.1%],F:0.4%,M:0.3%,n:2799 | 13790 |
| *Harpegnathos saltator* (Jerdon's jumping ant) | UP000008237 | 610380 | C:90.3%[S:90%,D:0.3%],F:6.4%,M:3.3%,n:4415 | 15029 |
| *Ooceraea biroi* (Clonal raider ant) | UP000053097 | 2015173 | C:95.9%[S:95.4%,D:0.4%],F:3.4%,M:0.8%,n:4415 | 16497 |
| *Papilio xuthus* (Asian swallowtail butterfly) | UP000053268 | 66420 | C:96.3%[S:94.9%,D:1.4%],F:2.4%,M:1.3%,n:1658 | 15265 |
| *Zootermopsis nevadensis* (Dampwood termite) | UP000027135 | 136037 | C:93.9%[S:92.8%,D:1.1%],F:1.6%,M:4.5%,n:1658 | 14539 |
| *Operophtera brumata* (winter moth) | UP000037510 | 104452 | C:75.9%[S:73.4%,D:2.5%],F:15%,M:9.2%,n:1658 | 16814 |
| *Lucilia cuprina* (Green bottle fly) | UP000037069 | 7375 | C:92.5%[S:91.5%,D:1%],F:1.8%,M:5.6%,n:2799 | 14353 |
| *Saccharomyces cerevisiae* (strain ATCC 204508 / S288c) (Baker's yeast) | UP000002311 | 559292 | C:98.9%[S:98.3%,D:0.6%],F:1.1%,M:0%,n:1711 | 6049 |
| *Emericella nidulans* (strain FGSC A4 / ATCC 38163 / CBS 112.46 / NRRL 194 / M139) (Aspergillus nidulans) | UP000000560 | 227321 | C:93%[S:92.9%,D:0.1%],F:4.5%,M:2.5%,n:4046 | 10557 |
| *Neurospora crassa* (strain ATCC 24698 / 74-OR23-1A / CBS 708.71 / DSM 1257 / FGSC 987) | UP000001805 | 367110 | C:99%[S:88.6%,D:10.4%],F:0.8%,M:0.2%,n:3725 | 9759 |
| *Yarrowia lipolytica* (strain CLIB 122 / E 150) (Yeast) (Candida lipolytica) | UP000001300 | 284591 | C:87.9%[S:87.3%,D:0.6%],F:9.8%,M:2.3%,n:1711 | 6449 |
| *Arachis hypogaea (*Peanut*)* | UP000289738 | 3818 | C:93.7%[S:17.2%,D:76.5%],F:1.7%,M:4.7%,n:1440 | 71122 |
| *Musa acuminata subsp. Malaccensis* (Wild banana) | UP000012960 | 214687 | C:86.7%[S:76.5%,D:10.3%],F:4.7%,M:8.5%,n:1440 | 36474 |
| *Arabidopsis thaliana* (Mouse-ear cress) | UP000006548 | 3702 | C:99.6%[S:59.1%,D:40.5%],F:0.2%,M:0.2%,n:1440 | 27466 |
| *Oryza sativa subsp. indica* (Rice) | UP000007015 | 39946 | C:94.8%[S:93.6%,D:1.2%],F:2.4%,M:2.8%,n:1440 | 37344 |
| *Zea mays* (Maize) | UP000007305 | 4577 | C:96.4%[S:49.4%,D:46.9%],F:2.1%,M:1.5%,n:1440 | 39400 |
| *Triticum aestivum* (Wheat) | UP000019116 | 4565 | C:99.4%[S:1.4%,D:98%],F:0.1%,M:0.5%,n:1440 | 105061 |
| *Physcomitrella patens subsp. patens* (Moss) | UP000006727 | 3218 | C:67.8%[S:52%,D:15.8%],F:3.3%,M:28.9%,n:1440 | 30857 |
| *Emiliania huxleyi* (Pontosphaera huxleyi) | UP000013827 | 2903 | C:74.9%[S:0.7%,D:74.3%],F:14.2%,M:10.9%,n:303 | 35676 |
| *Dictyostelium discoideum* (Slime mold) | UP000002195 | 44689 | C:96%[S:92.4%,D:3.6%],F:0.7%,M:3.3%,n:303 | 12739 |
| *Chlamydomonas reinhardtii* (Chlamydomonas smithii) | UP000006906 | 3055 | C:96%[S:90.1%,D:5.9%],F:2.3%,M:1.7%,n:303 | 17614 |
| *Thalassiosira pseudonana* (Marine diatom) | UP000001449 | 35128 | C:34.2%[S:33.3%,D:0.9%],F:1.3%,M:64.5%,n:234 | 11717 |
| *Plasmodium falciparum* (isolate 3D7) | UP000001450 | 36329 | C:25.6%[S:25.2%,D:0.4%],F:0.4%,M:73.9%,n:234 | 5376 |
